# G1-Cyclin2 (Cln2) promotes chromosome hypercondensation in *eco1/ctf7 rad61* null cells during hyperthermic stress in *Saccharomyces cerevisiae*

**DOI:** 10.1101/2022.05.27.493609

**Authors:** Sean Buskirk, Robert V. Skibbens

## Abstract

Eco1/Ctf7 is a highly conserved acetyltransferase that activates cohesin complexes and is critical for sister chromatid cohesion, chromosome condensation, DNA damage repair, nucleolar integrity, and gene transcription. Mutations in the human homolog of *ECO1* (*ESCO2/EFO2*), or in genes that encode cohesin subunits, result in severe developmental abnormalities and intellectual disabilities referred to as Roberts Syndrome (RBS) and Cornelia de Lange Syndrome (CdLS), respectively. In yeast, deletion of *ECO1* results in cell inviability. Co-deletion of *RAD61 (WAPL* in humans), however, produces viable yeast cells. These *eco1 rad61* double mutants, however, exhibit a severe temperature-sensitive growth defect, suggesting that Eco1 or cohesins respond to hyperthermic stress through a mechanism that occurs independent of Rad61. Here, we report that deletion of the G1 cyclin *CLN2* rescues the temperature sensitive lethality otherwise exhibited by *eco1 rad61* mutant cells, such that the triple mutant cells exhibit robust growth over a broad range of temperatures. While Cln1, Cln2 and Cln3 are functionally redundant G1 cyclins, neither *CLN1* nor *CLN3* deletions rescue the temperature-sensitive growth defects otherwise exhibited by *eco1 rad61* double mutants. We further provide evidence that *CLN2* deletion rescues hyperthermic growth defects independent of START and impacts the state of chromosome condensation. These findings reveal novel roles for Cln2 that are unique among the G1 cyclin family and appear critical for cohesin regulation during hyperthermic stress.

## INTRODUCTION

Cohesins (comprising Mcd1/Scc1, Smc1, Smc3, Irr1/Scc3, and Pds5 in budding yeast) are protein complexes that bind DNA to regulate a variety of essential cellular processes. During S phase, cohesins (dimers or clusters) tether together sister chromatids. This tethering, termed cohesion, is maintained until anaphase onset and thus ensures high fidelity chromosome segregation (Uhlmann and Nasmyth, 1998; Skibbens et al., 1999; Guacci et al., 1999; Michaelis et al., 1999; Cattoglio et al., 2019; Shi et al., 2020; Kulemzina et al., 2013; Tong and Skibbens, 2015; Eng et al., 2015; Zhang et al., 2008; Xiang and Koshland, 2021). In response to DNA damage during G2 and M phases, cohesins are recruited to the sites of double-strand breaks. This enhanced cohesion ensures template proximity, which in turn promotes homologous recombination and non-mutagenic DNA repair (Sjogren and Nasmyth 2001; Kim et al., 2002; Strom 2004; Unal 2004; Strom 2007; Unal 2007; Heidinger-Pauli 2008; Covo 2010; Lightfoot 2011; Kong 2014; Scherzer 2022; Piazza 2021; Mfarej 2020). During G1 phase, DNA-bound cohesins extrude DNA to form intramolecular loops. The length of the DNA loops in part are determined by the location of convergently oriented CTCF-bound DNA motifs, which halt cohesin-mediated DNA extrusion (Kim et al., 2019; de Wit et al., 2015; Davidson et al, 2016; Haarhuis 2017; Rao et al., 2017; Schwarzer et al., 2017; Gassler 2017; Wutz et al., 2017; Xiang and Koshland; Liu 2019). Numerous studies document that cohesin-dependent chromatin looping during G1 plays a critical role in both large- and small-scale conformations. Importantly, loops produced via cohesin-based DNA extrusion can bring into registration DNA regulatory elements (enhancers, insulators and promoters) that either induce or repress the transcription of individual genes (reviewed in Horsfield 2022; Mfarej and Skibbens 2020; Perea-Resa 2021). In this light, it is not surprising that mutations in cohesins, or cohesin regulators, result in genome-wide transcription dysregulation and severe birth defects (reviewed in Dorsestt 2016; Deardorff et al., 2005; Selicorni et al., 2021; Banerji et al., 2017). In concept, cohesin-based looping during G2/M could also promote chromosome condensation. While cohesin mutations indeed result in condensation defects, additional lines of evidence suggest that cohesins may promote condensation through condensin recruitment/activation (Guacci et al., 1999; Lavoie et al., 2002, 2004; Gard et al., 2009; Woodman et al., 2015; Ding 2006; Lamothe et al., 2020; Kakui and Uhlmann 2018; Skibbens 2019).

Eco1/Ctf7 (human ESCO2/EFO2 and ESCO1/EFO1) is a highly conserved acetyltransferase that activates cohesin via acetylation of either Smc3 during S phase or Mcd1 (human RAD21) during G2/M in response to DNA damage (Skibbens, Toth, Bellows, Williams et al., 2003; Heidinger-Pauli et al., 2009; Zhang et al., 2008; Rolef Ben-Shahar 2008; Unal et al., 2008). Eco1 is critical for sister chromatid cohesion, chromosome condensation, DNA damage repair, nucleolar integrity, and gene transcription (Skibbens et al., 1999; Strom et al., 2004; Unal et al., 2004; Strom et al., 2007; Unal et al., 2007; Heidinger-Pauli et al., 2008; Gard et al., 2009; Lu et al., 2010; Billon et al., 2017; Alomer et al., 2017; Wu et al., 2020; Zuilkoski and Skibbens 2020; Mfarej and Skibbens, 2020, 2022). Transcriptional dysregulations that arise in response to *ESCO2* mutation in humans result in severe developmental abnormalities and intellectual disabilities termed Roberts Syndrome (RBS) (Vega et al., 2005; Schule et al., 2005; Gordillo et al., 2008). Not surprisingly, RBS shares numerous phenotypes (Smithells and Newman, 1992) with those of Cornelia de Lange Syndrome (CdLS), which arises due to mutations in cohesin and other cohesin regulatory genes (Dorsestt 2016; Deardorff et al., 2005; Selicorni et al., 2021; Banerji et al., 2017). Numerous studies document the adverse effects of ESCO2/cohesin-dependent transcriptional dysregulation on tissue proliferation, bone development, immune function and promotion of tumorigenesis (Banerji et al., 2017; Banerji et al., 2017; Ketharnathan et al., 2020; Chin et al., 2020; Marsman et al., 2014; Rhodes et al., 2010; Horsfield et al., 2007; Monnich et al., 2011; Adane et al., 2021; Rogers et al., 2021). More recent findings now link RBS/CdLS genetic-based maladies to teratogenic reagents such as thalidomide (Sanchez et al., 2022). Thalidomide produces severe birth defects through the targeting of DDB1, a subunit of CRL4 ubiquitin ligase (Itoh et al., 2010). Studies in zebrafish embryos revealed that *ddb1* is transcriptionally co-regulated by both Esco2 and the cohesin subunit Smc3 and that exogenous *DDB1* expression rescues the developmental defects that arise due to SMC3 knockdown (Sanchez et al., 2022). Thus, the regulation of chromatin structure by Eco1/ESCO2 and cohesins during human development remains an issue of significant clinical relevance.

Eco1 is essential for cell viability in all model systems tested to date (Skibbens et al., 1999; Toth et al., 1999; Tanaki et al., 2000; Williams et al., 2003; Vega et al., 2005; Seitan et al., 2006; Kawauchi et al., 2009; Monnich et al., 2011; Whelan et al., 2012), with the exception of a clade of parasitic microbes that also lack cohesin-loading complexes (Scc2/NIPBL and Scc4/MAU-2) (Gentekaki et al., 2017) but which are otherwise similarly essential (Rollins et al., 1999; Ciosk et al., 2000; Tonkin et al., 2004; Krantz et al., 2004). One critical role of Eco1 is to establish cohesion between sister chromatids. To accomplish this, Eco1 is recruited to the DNA replication fork by PCNA and other fork associated components (Skibbens et al., 1999, Moldovan et al., 2006, Bender et al., 2020; Zhang et al., 2017; Zhang et al., 2018). At the fork, Eco1 acetylates the Smc3 subunit of cohesins that are newly deposited onto each sister chromatid.

Acetylation blocks cohesin dissociation from DNA (and possibly stabilizes cohesin dimerization or clustering), thereby establishing sister chromatid cohesion. Prior to acetylation, cohesins dissociate from DNA in a process that involves Rad61 (WAPL in humans), which binds the cohesin subunit Pds5 (Kueng et al., 2006; Ghandi et al., 2006; Sutani et al., 2009; Rowland et al., 2009). Cells devoid of Rad61 exhibit robust growth at all temperatures, similar to wildtype, even though they retain elevated levels of chromatin-bound cohesins. Intriguingly, cells that harbor deletions of both *ECO1* and *RAD61* survive (Sutani et al., 2008; Rolef Ben-Shahar et al., 2008), but only within a narrow temperature range (Maradeo et al., 2010; Guacci et al., 2012). The temperature sensitive lethality of *eco1 rad61* null cells suggest that Eco1 or cohesins perform an activity during hyperthermic stress that that is independent of Rad61.

Here, we report results from an unbiased genome-wide screen that identified the G1 cyclin Cln2 as an essential regulator of this Eco1/cohesin-dependent hyperthermic stress response. Our results reveal that deletion of *CLN2* rescues the temperature-dependent growth defects exhibited by *eco1 rad61* cells. While Cln1, Cln2 and Cln3 are functionally redundant in promoting the transition from G1 to S phase (START) (Lew et al., 1992; Lew et al., 1993; Levine et al., 1996; Queralt et al., 2004; Talarek et al., 2017), we find that Cln2 is unique among the G1 cyclins in rescuing *eco1* rad*61* double mutant cell growth defects. We further provide evidence that the suppression provided by *CLN2* deletion occurs independent of START and may act through alterations in chromatin structure.

## MATERIALS AND METHODS

### Yeast strains, media and growth conditions

All strains used in this study were derived from the S288C background and grown in YPD media unless otherwise noted (Supplemental Table 1). Diploid strains were sporulated in 0.3% potassium acetate and tetrads dissected on YPD agar. The genotypes of the resultant spores were analyzed for each wild type, single and double mutant spore recovered. Phenotypic analyses of isolated spores were performed by generating 10-fold serial dilutions of log phase cultures normalized based on OD_600_ as a measure of cell density. Each dilution series was plated on YPD agar plates, or selective medium plates, and grown at a range of temperatures as indicated in each figure. Non-ts revertant strains were tested for conversion to a ts-state after transformation with plasmid harboring either PDS5 (pVG177) or SCC3/IRR1 (pH9.3), kindly provided by Dr. Vincent Guacci (Stead et al., 2003), compared to vector alone. pVG177 and pH9 were independently verified by restriction digest. Primers used to delete genes and verify gene deletions are contained in Supplemental Table 2. Synchronization of yeast cultures and assessment of DNA contents by flow cytometry were performed as previously described (Maradeo and Skibbens, 2010; Tong and Skibbens, 2015).

Site-directed PCR mutagenesis was used to replace the entire *CLN2* open reading frame, or a portion of the *CLN2* reading frame (retaining sequences that encode for the first 221 residues), within the parental *eco1Δ rad61Δ* YMM828 strain (Maradeo and Skibbens, 2010) with a the *kanMX* selectable marker (Longtine et al., 1998). Appropriate knock-outs were confirmed by PCR (primer sequences are contained in Supplemental Table 2).

### Genomic sequencing

Genomic DNA was isolated from pools containing six individual segregants. Briefly, six segregants for each revertant were grown in YPD broth and cell densities normalized (based on OD_600_) prior to pooling. For each pooled sample, cell lysis was performed by physical agitation (Mini-Beadbeater, BioSpec) in the presence of 0.5mm glass beads (BioSpec) and lysis buffer (1% SDS, 100mM NaCl, 10mM Tris pH8.0; 2% Triton X-100) and Phenol:Chloroform:Isoamyl (25:24:1). Lysate was centrifuged (15,000 rpm, 5 min at 4°C), the aqueous supernatant incubated in RNAse (Sigma) and PCI re-extracted, genomic DNA obtained by centrifugation using Phase Lock Gel Light columns (Quanta), prior to precipitation by addition of 100% EtOH. DNA quality and concentration was assessed, following manufacturer’s instructions, using the Quant-it BR Assay kit for dsDNA (Invitrogen), prior to sending to the Microbial Genome Sequencing Center (Pittsburgh, PA) for sequencing on the Illumina NextSeq 500 instrument. FASTQ files were aligned to the S288C reference genome (Engel et al., 2013) using the Burrows-Wheeler Aligner with Maximal Exact Matches (BWA-MEM) algorithm version 0.7.15 with default settings (Li and Durbin, 2010). Variants were called using FreeBayes version 1.1.0 with default settings and the “--pooled-continuous” argument (Garrison and Marth, 2012). Calls were annotated using SnpEff version 4.3T with default settings (Cingolani et al., 2012). Mutations were manually verified by viewing in the Integrative Genomics Viewer (IGV) version 2.5 (Thorvaldsdóttir et al., 2013).

### Microscopy and condensation assay

Chromatin condensation assays were performed as previously described (Shen and Skibbens, 2017, 2020). DNA masses and rDNA structures were detected by DAPI staining. Chromatin images were captured using a Nikon Eclipse E800 microscope equipped with a cooled CD camera (Coolsnapfx, Photometrics) and IPLab software (Scanolytics). rDNA loops were often indistinguishable from chromatin masses in *eco1 rad61* double mutant cells. Attempts to include rDNA structural analyses resulted in the omission of a significant, if not majority, number of *eco1 rad61* double mutant cells. To accommodate the severity of this defect, we instead assessed genome-wide condensation. A region of interest was determined that matched the majority of wildtype genomic mass areas. Specifically, we experientially determined the size of a circular region of interest (ROI) or mask that was filled by wildtype DNA masses or in which the majority (approximately 3/4) of the edges coincided with wildtype DNA masses. The resulting mask (62 pixel circle) was then superimposed over each DNA mass to individually assess whether the DNA mass (excluding rDNA loop extensions) filled or failed to fill the ROI mask or contact the majority of the mask edges. This strategy enabled the inclusion of nearly 100% of all DNA masses (overlapping masses, in which genomic boundaries were obscured, were omitted from quantification).

### Flow cytometry and cell cycle progression

Cell synchronizations and cycle progression were monitored using propidium iodide and flow cytometry (BD FACScan) as previously described (Maradeo and Skibbens, 2010; Tong and Skibbens, 2015). Briefly, log phase cultures maintained at 30°C were normalized to an optical density typically between 0.1 and 0.4. For chromosome condensation assays, log phase cells were placed in fresh YPD supplemented with nocodazole (5 µg/ml final concentration) and maintained at 37°C for 3 hours prior to fixation. For cell cycle progression experiments, log phase cells were first synchronized in early S phase by maintaining growth at 30°C in fresh YPD supplemented with hydroxyurea (0.2M final concentration) before washing cells and resuspending them in fresh YPD supplemented with nocodazole (5 µg/ml final concentration). Cells were then shifted to 37°C and samples collected every 30 minutes. In either case, cells were spun and resuspended in fixing solution (0.2M Tris 70% EtOH solution). Cells were then treated with RNase (Roche) and proteinase K (Roche) solutions to remove RNA and protein, respectively. To analyze DNA content, cells were stained with a 0.0001% propidium iodide (Sigma) solution. Prior to use, the stock solution was diluted 10ul + 990ul in 0.2M Tris solution for each milliliter of sample. Cells were sonicated and DNA content quantified by flow cytometry using a BD FACSCanto II.

### Statistical analyses

A chi-square test of independence was used to assess the relationship of gene mutations (*eco1Δ rad61Δ* double mutant cells and *eco1Δ rad61Δ cln2Δ* triple mutant cells) on chromatin condensation levels, standardized using wildtype cells. For each strain, the number of observed genomic masses that matched or exceeded the template were compared to expected (all genomic masses in the field of view) number of genomic masses. A statistical significance p-value at or below 0.05 was used to demonstrate gene mutation dependence. We also compared chromatin condensation effects across the three strains. Here, the percentage of genomic masses, within a given field, that matched or exceeded the template were used for subsequent analysis, performed using a two-tailed Student’s t-tests to assess statistical significance using a p-value at or below 0.05 between each group.

### Data availability statement

Strains and plasmids are available upon request. The authors affirm that all data necessary for confirming the conclusions of the article are present within the article, figures, and tables. The short read sequencing data have been deposited in the NCBI BioProject database (PRJNA836598).

## RESULTS

### A spontaneous suppressor screen identifies potential regulators of Eco1 function

Budding yeast Eco1/Ctf7 (Eso1 in *S. pombe*, ESCO1,2 in vertebrates) is the founding member of a highly conserved family of acetyltransferases that are essential regulators of cohesin function (Skibbens et al., 1999; Tanaka et al., 2000; Ivanov et al., 2002; Zhang et al., 2008; Rolef Ben-Shahar 2008; Unal et al., 2008). While mutations in a small number of cohesin pathway genes (*PDS5*, *SCC3/IRR1* and *RAD61*) were initially reported to bypass Eco1 function, the resulting double mutant cells (for instance, *eco1Δ rad61Δ* double mutated cells) are viable within only a narrow temperature range (Sutani et al., 2009; Rowland et al., 2009; Maradeo and Skibbens, 2010; Guacci and Koshland, 2012). These findings suggest that Eco1 and cohesin perform essential functions in response to hyperthermic stress that are independent of Rad61. We focused on the suppression provided by *RAD61* deletion since Rad61 is not a core cohesin component and *RAD61* loss is sufficient to impact chromatin structure in the absence of other cohesin subunit mutations (Kueng et al., 2006; Ghandi et al., 2006).

Prior to performing a genome-wide suppressor screen, we first independently documented the hyperthermic sensitivity of *eco1Δ rad61Δ* double mutant cells. Similar to wildtype cells, *rad61Δ* single mutant cells exhibited robust growth at all temperatures tested (Figure 1A). In contrast, *eco1^ctf7-203^* mutant cells exhibited severe growth defects at 30°C and were inviable at 37°C (Figure 1A) (Skibbens et al., 1999). *eco1Δ rad61Δ* double mutant cells exhibited robust growth at 30°C but were inviable at 37°C (Figure 1A), in support of earlier findings that *eco1Δ rad61Δ* cells exhibit severe hypothermic growth sensitivity (Sutani et al., 2009; Maradeo and Skibbens, 2010; Guacci and Koshland, 2012).

**Figure 1.**
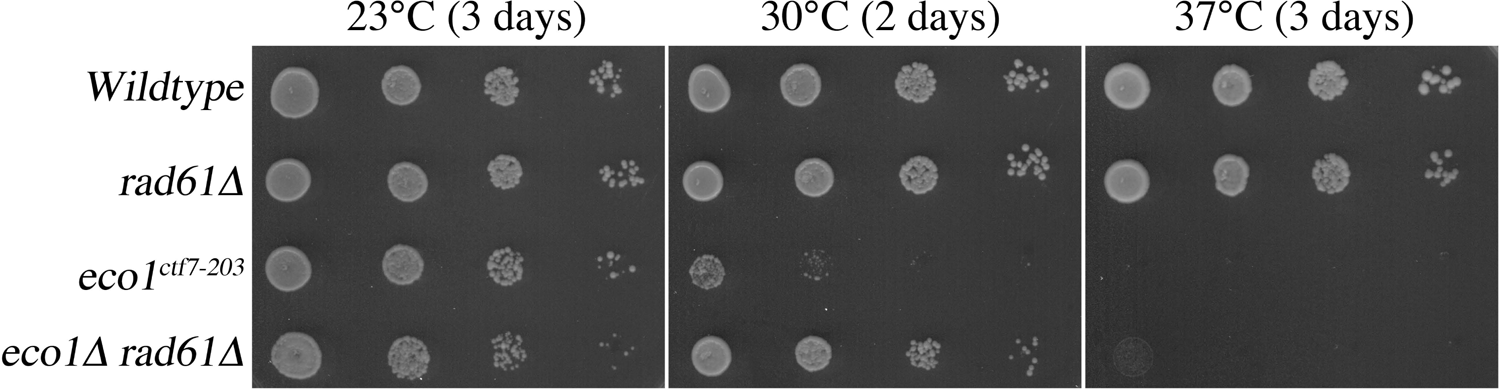
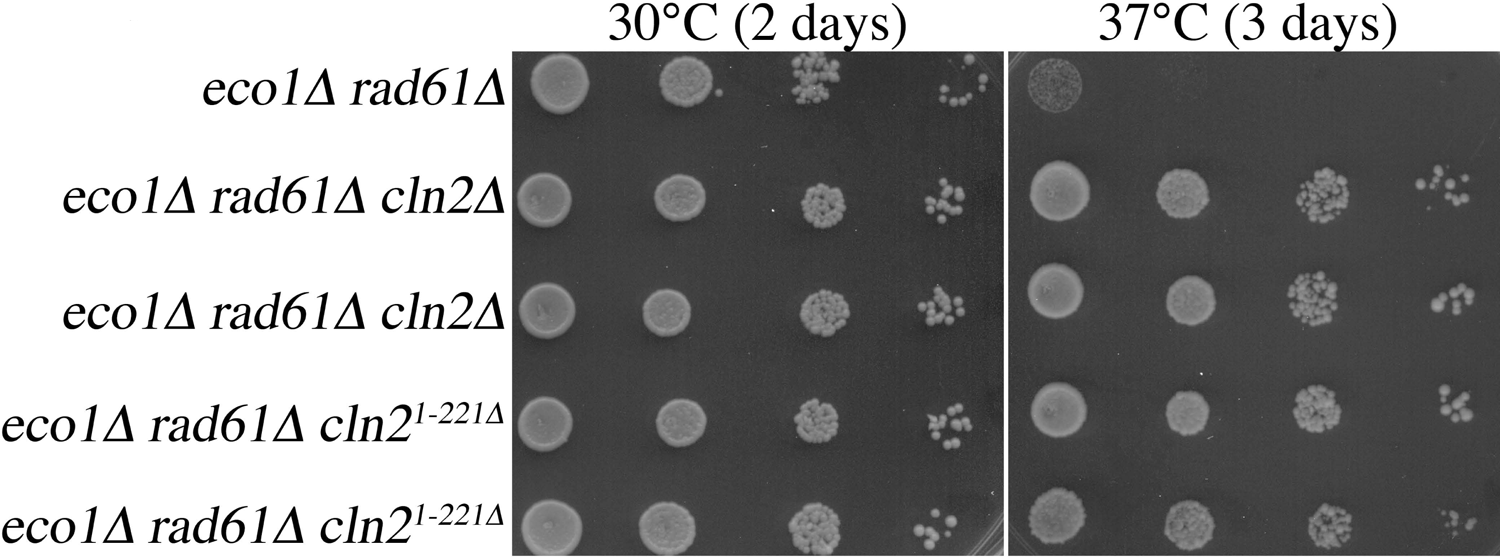
Identification of the G1 cyclin, Cln2, as a regulator of Eco1-Rad61 pathways. A) *eco1Δ rad61Δ* double mutant cells are viable within a narrow temperature range. 10-fold serial dilutions of wildtype, *rad61Δ* and *eco1^ctf7-203^* single mutant cells, and *eco1Δ rad61Δ* double mutant cells at the indicated temperatures. B) *CLN2* deletion, and truncated Cln2^1-221^ (*CLN2^1-221Δ^*), both suppress *eco1Δ rad61Δ* double mutant cells growth defects. 10-fold serial dilutions of wildtype cells, two independent isolates of *eco1Δ rad61Δ cln2Δ* triple mutant cells, and two independent isolates of *eco1Δ rad61Δ cln2^1-221^* cells grown at the indicated temperatures.

To identify novel suppressors of *eco1Δ rad61Δ* cell hyperthermic growth defects, two independently derived *eco1Δ rad61Δ* double mutant cells, YMM828 and YMM829 (Maradeo and Skibbens, 2010), were grown to log phase before plating 5×10^7^ cells onto YPD medium. After 4 days growth at 37°C, the plates were replica-plated onto YPD agar and incubated at 37°C for an additional two days. From each plate, we isolated ten moderate-to-large sized colonies, each containing spontaneous mutations that support growth under hyperthermic stress. From here on, these 20 strains will be referred to as revertants since each spontaneous mutation reverts temperature sensitive (ts) growth to a non-ts growth status. Since our goal was to identify novel suppressors of *eco1* mutant cell defects, each of the 20 triple-mutant revertants (*eco1Δ rad61Δ* and the reverting mutated gene) was transformed with plasmid alone or plasmid harboring either *PDS5* or *SCC3/IRR1*. Resulting *PDS5* or *SCC3/IRR1* transformants that exhibited cell inviability at 37°C were excluded from further analyses.

Spontaneous gene mutations that rescue *eco1Δ rad61Δ* mutant cell ts growth defects likely arise concordantly with other gene mutations that are unrelated to the revertant phenotype. To reduce the number of unrelated mutations, the remaining revertants were back-crossed two times to wildtype cells. In each case, the reverting mutation segregated according to Mendelian genetics (data not shown), suggesting that a single gene mutation was responsible for the revertant phenotype. To aid in the identification of the suppressor mutations, multiple segregants, derived from the same revertant, were pooled together based on the logic that only the suppressor mutation would be present in all segregants (present at a frequency of 1.0 in the pool). In contrast, non-revertant mutations were expected to segregate randomly and thus be present at intermediate or low frequencies. After pooling the segregants for each revertant (revertant A and B), genomic DNA was harvested and whole genome sequencing performed. Each of the two segregant pools (Pool A and B) contained two mutations at a frequency of 1.0 (Table 1). Of the two mutations (*CLN2* and *SMC2* for Pool A, *CLN2* and *RPL31A* for Pool B), only the *CLN2* mutation (frameshift at threonine 222), was present at a frequency of 1.0 in both pools (Table 1), suggesting that this allele might be the suppressor mutation and arose independently in these revertant strains.

**Table 1.** Identification of spontaneous revertant mutations in Revertant Pools A and B.

| Pool A |  |  |  |  |  |
| --- | --- | --- | --- | --- | --- |
| Chromosome Position | DNA Change | Annotation | Gene | Description ^ | Frequency in Pool |
| chrXVI_65946 | <u>delACTT</u> | frameshift at T222 | *CLN2 (YPL256C)* | G1 cyclin involved in regulation of the cell cycle | 1.00 |
| chrVI_218888 | A>G | K407E | *SMC2 (YFR031C)* | Subunit of the <u>condensin</u> complex | 1.00 |
| chrIII_297908 | G>C | S233T | *GIT1 (YCR098C)* | Plasma membrane permease | 0.72 |
| chrXIV_90717 | G>C | V139L | *CAF40 (YNL288W)* | Component of the CCR4-NOT transcriptional complex | 0.66 |
| chrII_98352 | G>C | L588F | *SEF1 (YBL066C)* | Putative transcription factor | 0.49 |
| chrVIII_484285 | <u>delT</u> | Y87fs | *PTH1 (YHR189W)* | One of two mitochondrially-localized peptidyl-tRNA hydrolases | 0.47 |
| chrXVI_252349 | A>T | E654D | *AIM44 (YPL158C)* | Regulator of Cdc42p and Rho1p | 0.47 |
| chrIV_178744 | G>T | W137C | *MSH5 (YDL154W)* | Protein of the <u>MutS</u> family | 0.37 |
| chrIV_322263 | C>T | T13I | *RPL31A (YDL075W)* | Ribosomal 60S subunit protein L31A | 0.25 |
| Pool B |  |  |  |  |  |
| Chromosome Position | DNA Change | Annotation | Gene | Description ^ | Frequency in Pool |
| chrXVI_65946 | <u>delACTT</u> | frameshift at T222 | *CLN2 (YPL256C)* | G1 cyclin involved in regulation of the cell cycle | 1.00 |
| chrIV_322263 | C>T | T13I | *RPL31A (YDL075W)* | Ribosomal 60S subunit protein L31A | 1.00 |
| chrIV_1193539 | T>C | I449T | *EAF1 (YDR359C)* | Component of the NuA4 histone acetyltransferase complex | 0.43 |
| chrXVI_133275 | C>G | A78G | *FLC1 (YPL221W)* | Flavin adenine dinucleotide transporter | 0.31 |
| chrV_191025 | C>T | A255V | *AFG3 (YER017C)* | Mitochondrial inner membrane m-AAA protease component | 0.27 |

^ from SGD

### *CLN2* deletion suppresses *eco1Δ rad61Δ* cell growth defects

Despite the high frequency of the *CLN2* mutation in all segregants derived from the two revertants, it became important to independently confirm that the *CLN2* mutation is responsible for the revertant phenotype. In the first of two strategies, the complete coding sequence of *CLN2* was deleted from our parental *eco1Δ rad61Δ* strain. In the second strategy, we constructed a partial deletion of *CLN2* (in which only the first 221 amino acids of Cln2 are retained in the genome), to replicate the allele obtained in our spontaneous suppressor screen (Table 1). In both cases, site-directed PCR mutagenesis of *CLN2* was confirmed by PCR in the resulting transformants (Longtine et al., 1998). Log phase cultures of the parental *eco1Δ rad61Δ* double mutant strain, and two independent transformants each of *eco1Δ rad61Δ cln2Δ* and *eco1Δ rad61Δ cln2^1−221^* triple mutant strains, were serially diluted, plated onto YPD agar, and incubated at either 30°C or 37°C. As expected, all strains exhibited robust growth at 30°C while *eco1Δ rad61Δ* double mutant cells were inviable at 37°C. Both isolates of the *eco1Δ rad61Δ cln2Δ* triple mutant cells, as well as the *eco1Δ rad61Δ cln2^1−221^* triple mutant cells, exhibited substantial growth at 37°C (Figure 1B). These results confirm that *CLN2* loss-of-function mutations rescue the ts growth defect otherwise present in *eco1Δ rad61Δ* double mutant cells.

### Cln2 is unique among G1 cyclins in suppressing *eco1Δ rad61Δ* cell growth defects

Cln1, Cln2, and Cln3 are functionally redundant such that any single G1 cyclin, bound to cyclin-dependent kinase (CDK), can promote START. Cln3-CDK, however, is the earliest inactivator of Whi5, a key inhibitor of START, that acts in both larger mother cells and smaller newly budded cells (reviewed Fisher, 2016; Ewald, 2018; Li et al., 2021). We thus tested whether deletion of *CLN3*, using site-directed PCR, would suppress *eco1Δ rad61Δ* double mutant cell ts growth defects, similar to deletion of *CLN2*. As expected, *eco1Δ rad61Δ* double mutant cells and *eco1Δ rad61Δ cln3Δ* triple mutant cells (three isolates shown) exhibited robust growth at 30°C. Surprisingly, and unlike *eco1Δ rad61Δ cln2Δ* cells, all isolates of *eco1Δ rad61Δ cln3Δ* triple mutant cells were inviable at 37°C and provided no growth advantage at 23°C (Figure 2A). Thus, *CLN3* deletion does not phenocopy the revertant growth defect provided by *CLN2* deletion during hyperthermic stress, revealing a novel distinction between these G1 cyclins.

**Figure 2.**
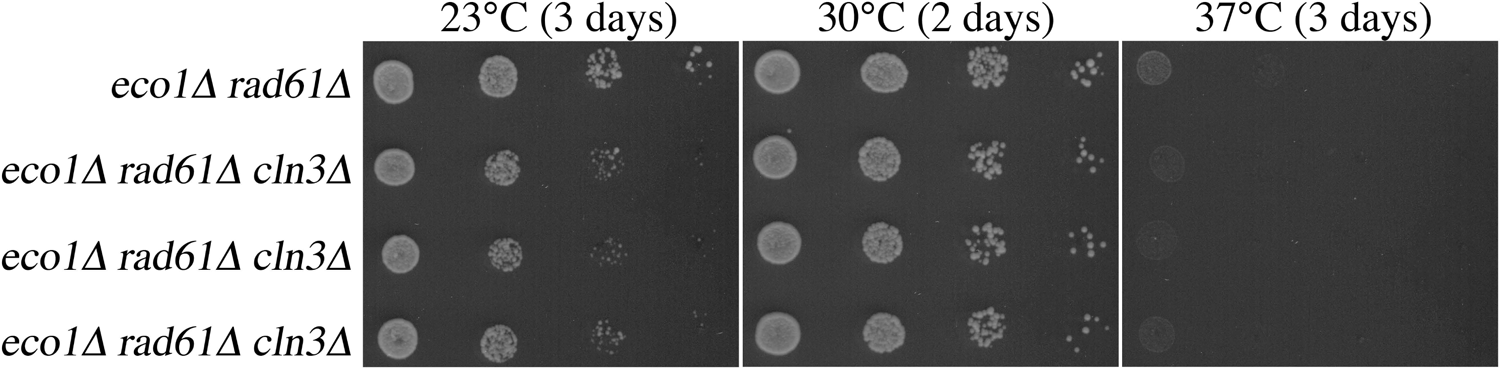
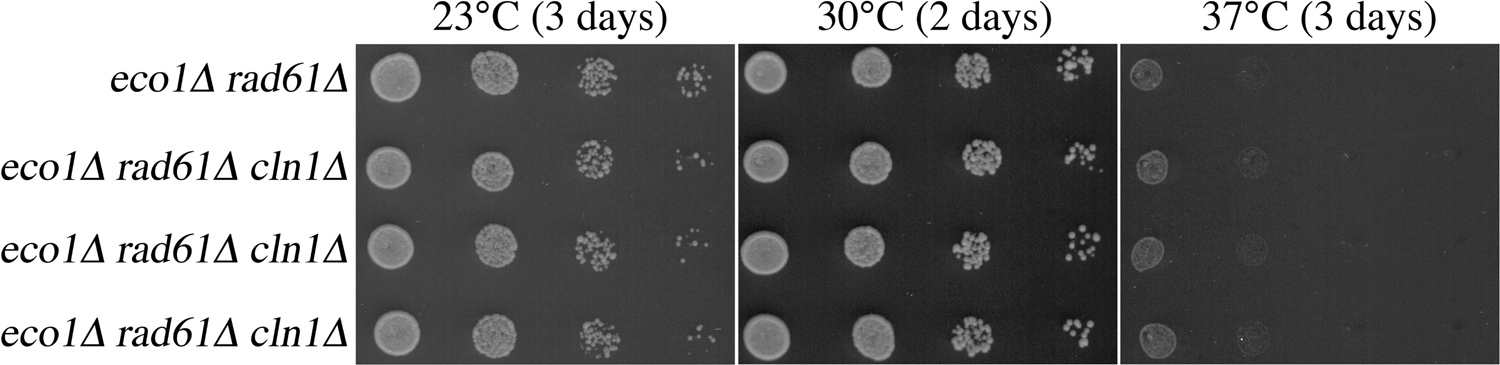
The role of Cln2 in Eco1-Rad61 functions is unique from Cln1 and Cln3 G1 cyclins. A) *CLN3* deletion does not rescue *eco1Δ rad61Δ* cell growth defects under thermic stress. 10-fold serial dilutions of parental *eco1Δ rad61Δ* double mutant cells and 3 independent isolates of *eco1Δ rad61Δ cln3Δ* triple mutant cells grown at the indicated temperatures. B) *CLN1* deletion does not rescue *eco1Δ rad61Δ* cell growth defects under thermic stress. 10-fold serial dilutions of parental *eco1Δ rad61Δ* double mutant cells and 3 independent isolates of *eco1Δ rad61Δ cln1Δ* triple mutant cells grown at the indicated temperatures.

Cln1 and Cln2 are roughly 60% identical at the amino acid level and bind identical target-docking sites. Cln1-CDK and Cln2-CDK are also functionally redundant in regulating Whi5 inactivation in smaller bud cells that arise due to the asymmetric division in budding yeast. Given the highly conserved roles of Cln1 and Cln2, we predicted that the deletion of *CLN1* would suppress *eco1Δ rad61Δ* double mutant cell ts growth defects, similar to the deletion of *CLN2*. We used site-directed PCR to generate *eco1Δ rad61Δ cln1Δ* triple mutant cells and assessed these for growth at 23°C and 30°C and the revertant phenotype at 37°C. Under all conditions, growth of *eco1Δ rad61Δ cln1Δ* triple mutant matched that of *eco1Δ rad61Δ* double mutant: each strain grew at 30°C but were inviable at 37° (Figure 2B). In combination, these results reveal that Cln2-CDK performs an activity that is distinct from both Cln1-CDK and Cln3-CDK.

### Cln2 antagonizes Eco1 functions independent of Rad61

*RAD61* deletion does not fully bypass the essential function of Eco1, such that *eco1 rad61* cells are inviable at 37°C (Figure 1) (Maradeo and Skibbens, 2010; Guacci and Koshland, 2012). Given that *CLN2* deletion fully rescues *eco1Δ rad61Δ* cell growth under hyperthermic conditions, it became important to test whether deletion of *CLN2* would fully bypass Eco1 function or only partially suppress Eco1 function, similar to deletion of *RAD61*. To differentiate between these two possibilities, *CLN2* was deleted from *eco1^ctf7-203^* ts cells (Skibbens et al., 1999). Log phase cultures of wildtype cells, *eco1^ctf7-203^* single mutant cells, and *eco1^ctf7-203^ cln2Δ* double mutant strains were serially diluted, plated onto YPD plates, and maintained at either 23°C, 30°C, or 37°C. As expected, all strains exhibited robust growth at 23°C (Figure 3). Consistent with prior studies, *eco1^ctf7-203^* single mutant cells exhibited severe ts growth defects at 30°C and were inviable at 37°C. *eco1^ctf7-203^ cln2Δ* double mutant strains (three isolates shown) exhibited significant improvement in growth at 30°C, compared to *eco1^ctf7-203^* single mutant cells but were inviable at 37°C (Figure 3). Our finding that the individual deletion of either *RAD61* or *CLN2* deletion fails to rescue *eco1* mutant cell ts growth (Figures 1 and 2), but that the combined deletion of *RAD61* and *CLN2* provides for robust growth at elevated temperatures, suggests that each exhibits a distinct activity on cohesins or Eco1-dependent regulation of cohesins (see Discussion).

**Figure 3.**
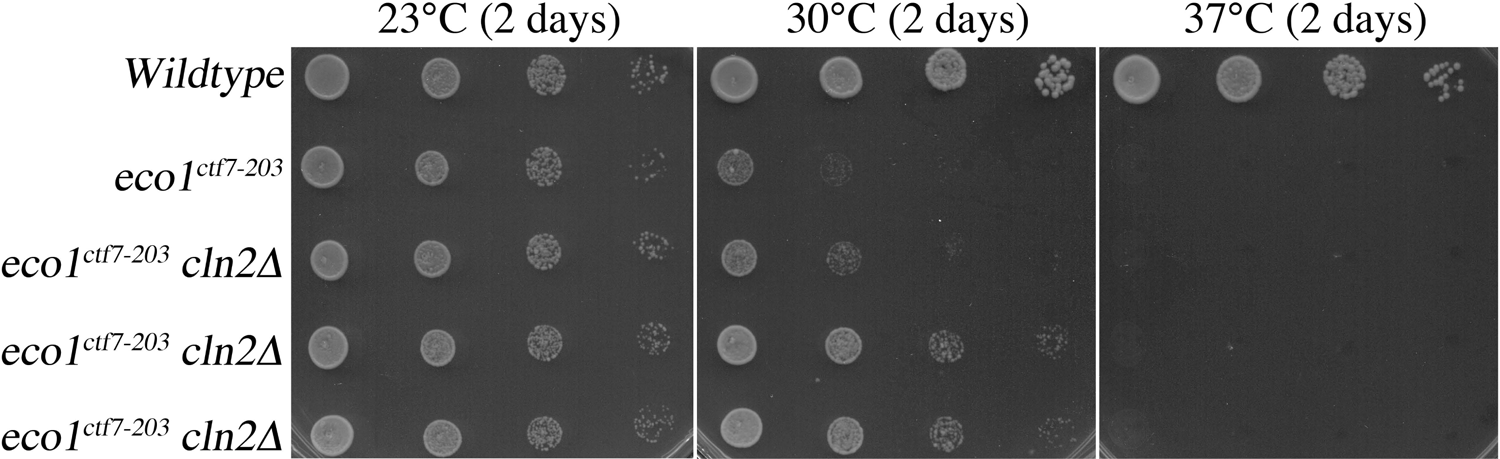
Deletion of *CLN2* suppresses *eco1* mutant cell ts growth, similar to but distinct from deletion of *RAD61*. 10-fold serial dilutions of wildtype cells, *eco1^ctf7-203^* mutant cells, and 3 independent isolates of *eco1^ctf7-203^ cln2Δ* double mutant cells grown at the indicated temperatures.

### *CLN2* deletion-dependent suppression of *eco1Δ rad61Δ* cell growth defects appears distinct from coupling cohesion establishment to DNA replication

The partial suppression of *eco1Δ* cell ts growth defects by mutation of either *CLN2*, *PDS5,* or *SCC3/IRR1* (Sutani et al., 2009; Rowland et al., 2009; Rolef Ben-Shahar et al., 2008; Maradeo and Skibbens, 2010; current study) likely arises due to a coordinated reduction of opposing activities that rebalance some inherent cohesin activity. In contrast, overexpression of wildtype genes that suppress *eco1* mutant cell growth defects are more likely gene products that augment Eco1 function. For instance, elevated levels of PCNA (*POL30*) suppresses *eco1^ctf7-203^* ts growth defects (Skibbens et al., 1999), a finding that revealed that the tethering together of sister chromatids is intimately linked to DNA replication. It thus became important to ascertain whether *eco1Δ rad61Δ* cell growth defects might be suppressed by PCNA overexpression, possibly linking the effects of the *CLN2* deletion to a DNA replication-based mechanism. To test this possibility, we overexpressed PCNA from a high-copy plasmid that harbors *POL30* (Skibbens et al., 1999). *eco1Δ rad61Δ* double mutant cells, harboring vector alone or vector driving elevated PCNA levels, exhibited strong growth at 30°C but were inviable when maintained at 37°C (Figure 4). Thus, the suppression that results from *CLN2* deletion is likely independent of the beneficial effect of coupling cohesion establishment to DNA replication.

**Figure 4.**
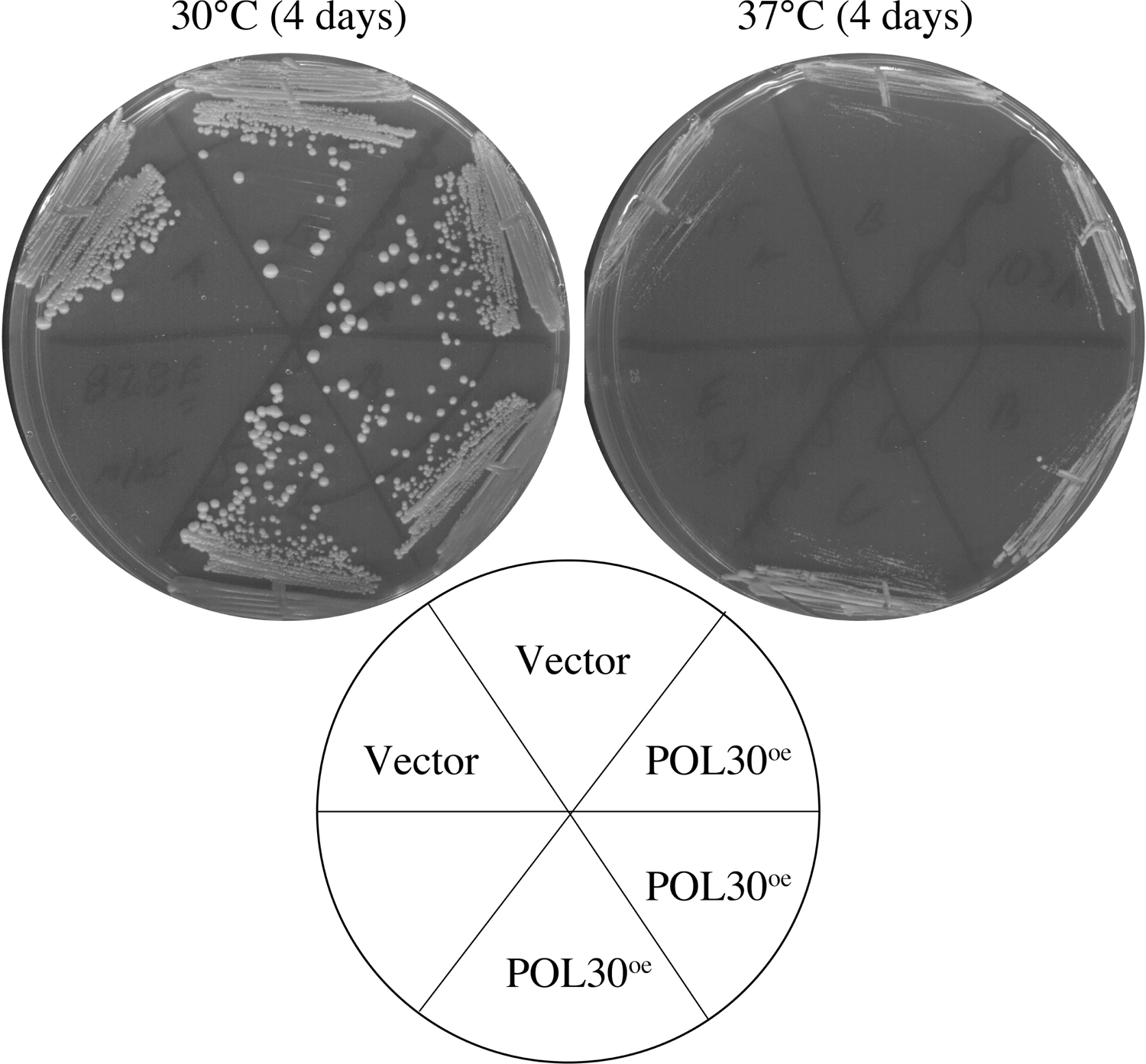
Elevated expression of PCNA (*POL30*) does not suppress *eco1Δ rad61Δ* mutant cell growth defects during thermic stress. The growth, two temperatures, of two independent isolates of *eco1Δ rad61Δ* double mutant cells transformed with vector alone compared to 3 independent isolates of *eco1Δ rad61Δ* double mutant cells transformed with vector that directs elevated *POL30* expression.

### *cln2Δ-dependent* suppression of *eco1Δ rad61Δ* cell growth defects occurs independent from a delay in START

Given the role of Cln2-CDK in promoting the transition from G1 to S phase, we wondered whether the *cln2Δ*-dependent suppression of *eco1Δ rad61Δ* cell growth defects might be due to slowed cell cycle progression. We tested this possibility in the following two ways. First, log phase cultures of wildtype cells and two independent isolates each of *eco1Δ rad61Δ* double mutant cells and *eco1Δ rad61Δ cln2Δ* triple mutant cells were grown at 30°C (permissive for all five strains) prior to shifting to 37°C. The DNA content in each condition was quantified by flow cytometry (Figure 5). As expected, wildtype cells exhibited log phase growth characteristics (indicated by roughly equivalent G1 and G2/M DNA peaks), such that pre- and post-replication peaks of DNA were unaltered even after prolonged growth at 37°C (Figure 5). Independent isolates of *eco1Δ rad61Δ* double mutant cells exhibited a G2/M bias in their DNA profiles both at 30°C and 37°C. This cell cycle delay is consistent with the cohesion and condensation defects that persist in these cells and that likely trigger a mitotic checkpoint (Guacci et al., 2012). In contrast, the DNA profiles of *eco1Δ rad61Δ cln2Δ* triple mutant cells appeared unaffected by growth at 37°C, exhibiting similar G1 and G2/M peaks as wildtype cells (Figure 5). Thus, loss of *CLN2* does not result in the accumulation of cells in a G1/S transition state but instead appears to promote cell cycle progression.

**Figure 5.**
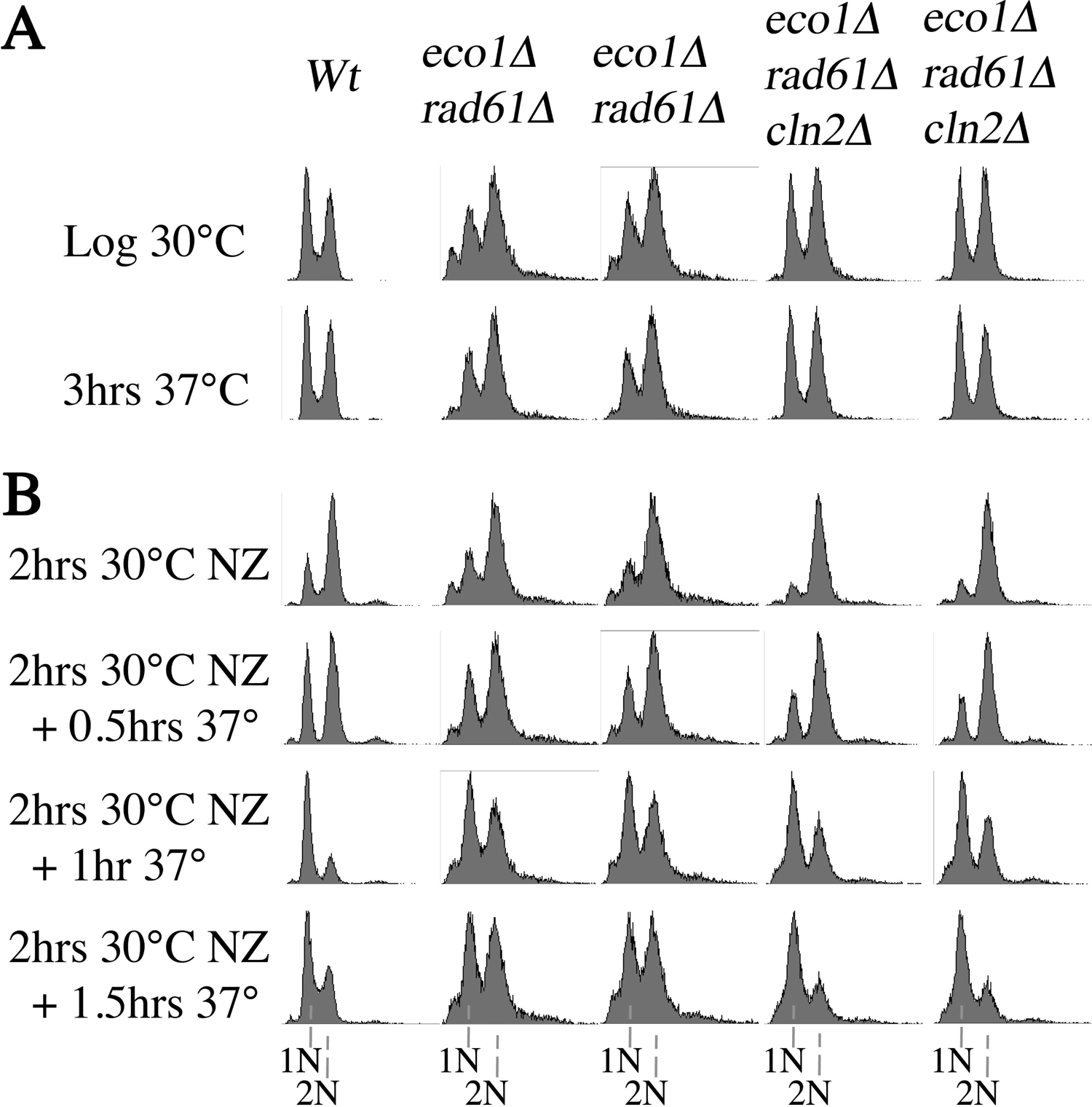
Deletion of *CLN2* does not delay, but instead promotes, cell cycle progression in *eco1Δ rad61Δ* mutant cells. A) DNA contents assessed using flow cytometry of log phase wildtype cells, 2 independent isolates of parental *eco1Δ rad61Δ* double mutant cells, and 2 independent isolates of *eco1Δ rad61Δ cln2Δ* triple mutant before and after growth at 37°C for 3 hours. B) DNA contents of the cells, described in (A), released from a mitotic arrest (NZ, nocodazole) at the permissive temperature of 30°C then shifted to 37°C. Samples harvested every 30 minutes were analyzed for DNA content by flow cytometry.

We next mapped cell cycle progression of wildtype cells, *eco1Δ rad61Δ* double mutant cells, and *eco1Δ rad61Δ cln2Δ* triple mutant cells at greater temporal resolution. Log phase cultures of each were arrested in preanaphase at 30°C, a temperature permissive for all of the strains, by the addition of nocodazole into the medium. After two hours, cells were washed free of nocodazole, incubated at 37°C in fresh medium and samples harvested every 30 minutes for analysis by flow cytometry. Wildtype cells exhibited a strong G2/M (post-replication) DNA content in nocodazole-arrested cells, but within the first 30 minute-time point, approximately half the cells exited mitosis and contained a G1 content of DNA. Within the first hour, almost all wildtype cells had transitioned to a G1 (pre-DNA replication) state (Figure 5). In contrast, neither isolate of *eco1Δ rad61Δ* double mutant cells exhibited significant cell progression at either the 30 minute or 1 hour time point (Figure 5). The *eco1Δ rad61Δ cln2Δ* triple mutant cells exhibited a slight delay in cell cycle progression, compared to the wildtype cells but far outpaced the progress of *eco1Δ rad61Δ* double mutant cells. Thus, loss of Cln2 promotes cell cycle progression cells, suggesting that the Cln2-CDK activity that antagonizes Eco1 occurs independent of START.

### *CLN2* deletion suppresses the hypercondensation defect in *eco1Δ rad61Δ* double mutant cells

The positive effect of *CLN2* deletion appears to be independent of both START regulation (Figure 5) and the coupling of cohesion establishment to DNA replication (Figure 4). *ECO1* mutation not only abrogates the establishment of cohesion during S phase, but also produces condensation defects of mitotic chromosomes (Skibbens et al., 1999; Gard et al., 2009; Zuilkoski and Skibbens, 2020). Rad61/WAPL is another regulator of chromatin condensation, but loss of Rad61/WAPL results in hypercondensed chromosomes (Lopes-Serra et al., 2013; Tedeschi et al., 2013; Wutz et al., 2017). Since both Eco1 and Rad61 impact condensation, we tested the extent to which *CLN2* deletion impacted chromosome condensation in *eco1Δ rad61Δ* double mutant cells. Notably, even wildtype cells respond to hyperthermic stress by increased compaction along the rDNA locus (Shen and Skibbens, 2017; Matos-Perdomo and Machin, 2018). The rDNA locus, which forms a puff-like structure during G1 but condenses into a tight loop or linear structure during mitosis, is a prominent feature analyzed in numerous condensation studies (Guacci et al., 1997; Sullivan et al., 2004; D’Ambrosio et al., 2008; Lopez-Serra et al., 2013; de Los Santos-Velázquez et al., 2017; Shen and Skibbens, 2007; Matos-Perdermo et al., 2018; Srinivasan et al., 2018; Lamothe and Koshland, 2020). Thus, we performed a condensation assay, modified from FISH protocols, that provides for high-resolution analyses of both genomic mass areas and rDNA structures in response to elevated temperatures (Guacci et al., 1994; Guacci et al., 1997; Shen and Skibbens, 2017; Shen and Skibbens, 2020; Srinivasan et al., 2018; Boginya et al., 2019).

To first assess the effect of *CLN2* deletion on chromatin condensation, log phase cultures of wildtype cells, *eco1Δ rad61Δ* double mutant cells, and two independent isolates of *eco1Δ rad61Δ cln2Δ* triple mutant cells, were released into fresh medium supplemented with nocodazole and incubated at 37°C for 3 hours. Cell cycle arrest was monitored by flow cytometry (Figure 6A) and chromatin condensation assessed by fluorescence microscopy as previously described (Shen and Skibbens, 2017). As expected, preanaphase wildtype cells contained easily discernible rDNA that formed discrete loops that emanated from the bulk of the genomic mass (Figure 6B). In *eco1Δ rad61Δ* double mutant cells, however, rDNA loops appeared reduced in length (Figure 6B), but most often were indiscernible such that attempts to analyze rDNA structures resulted in the exclusion of a significant population of cells. Despite this, easily discernible rDNA loops were clearly evident in *eco1Δ rad61Δ cln2Δ* triple mutant cells, qualitatively suggesting that *CLN2* deletion rescue the hypercondensation defects otherwise observed in *eco1Δ rad61Δ* double mutant cells.

**Figure 6.**
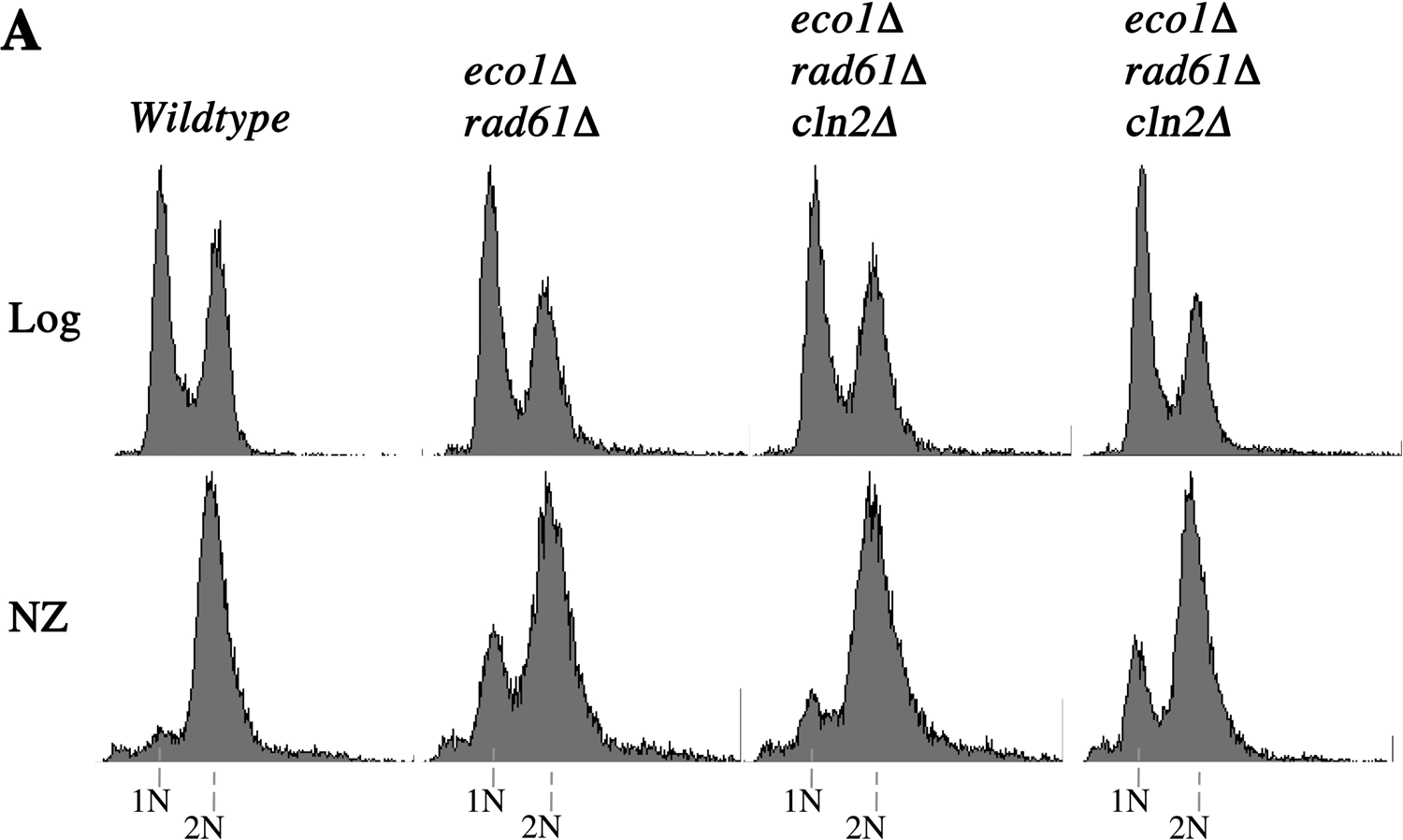
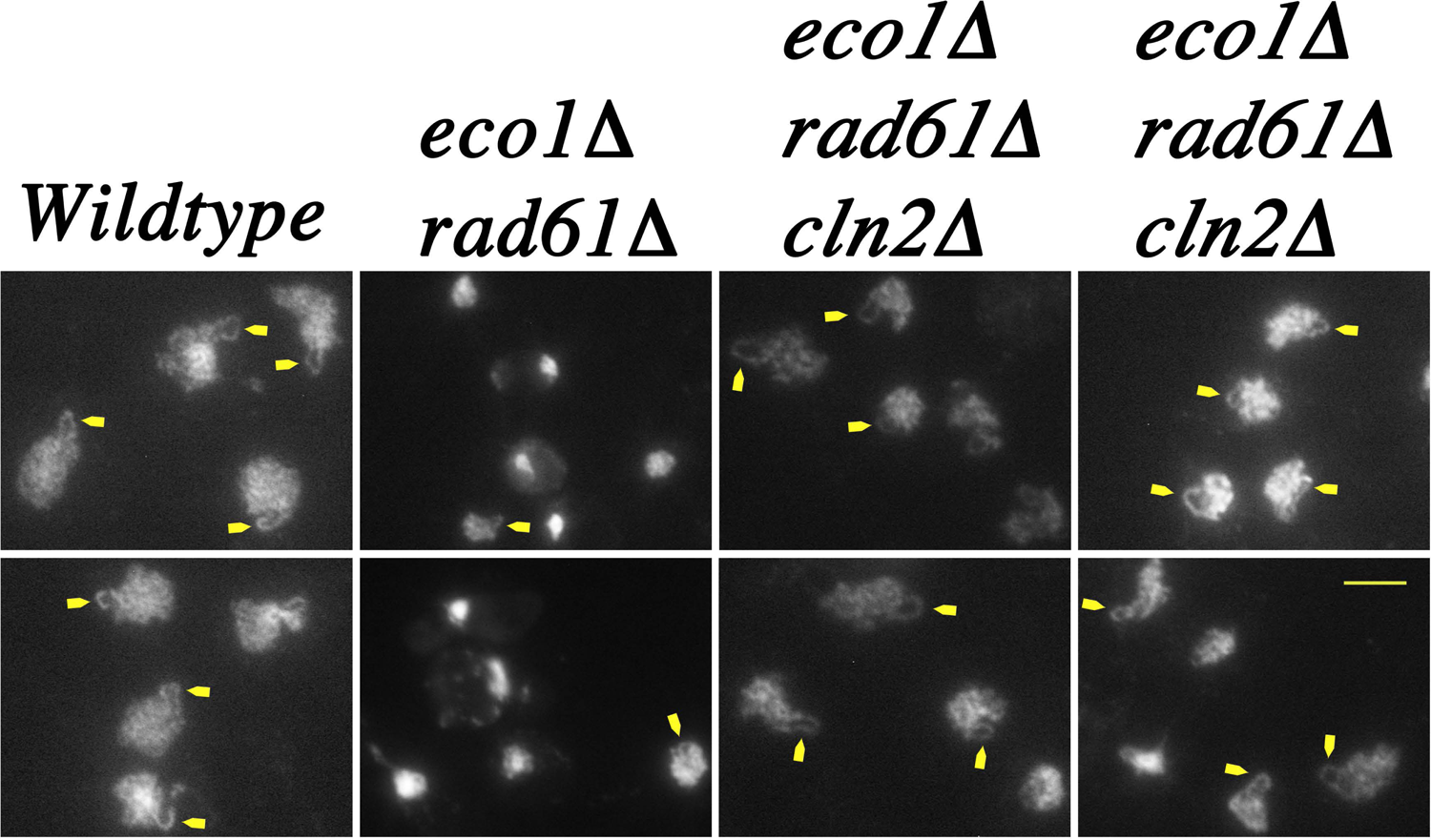
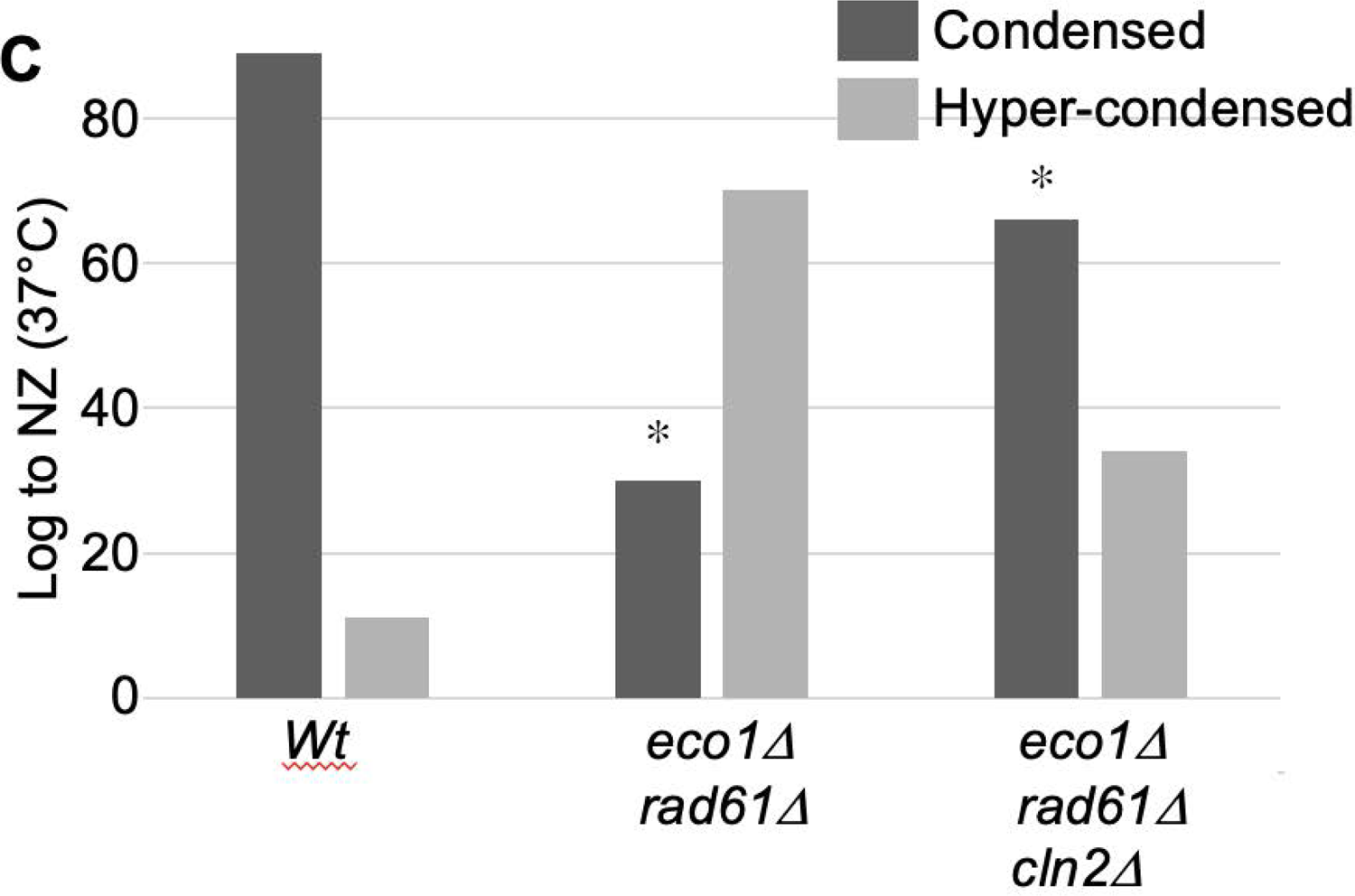
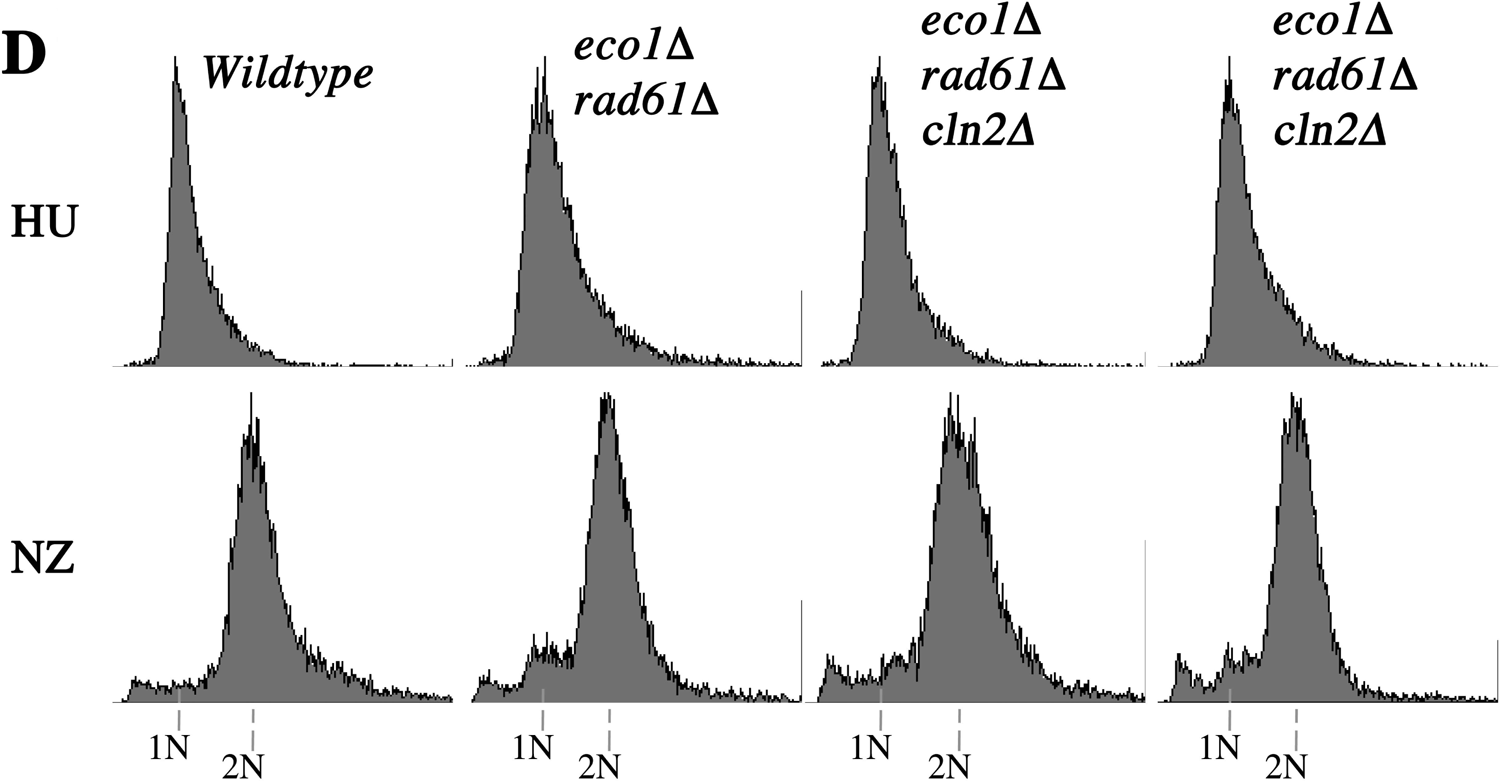
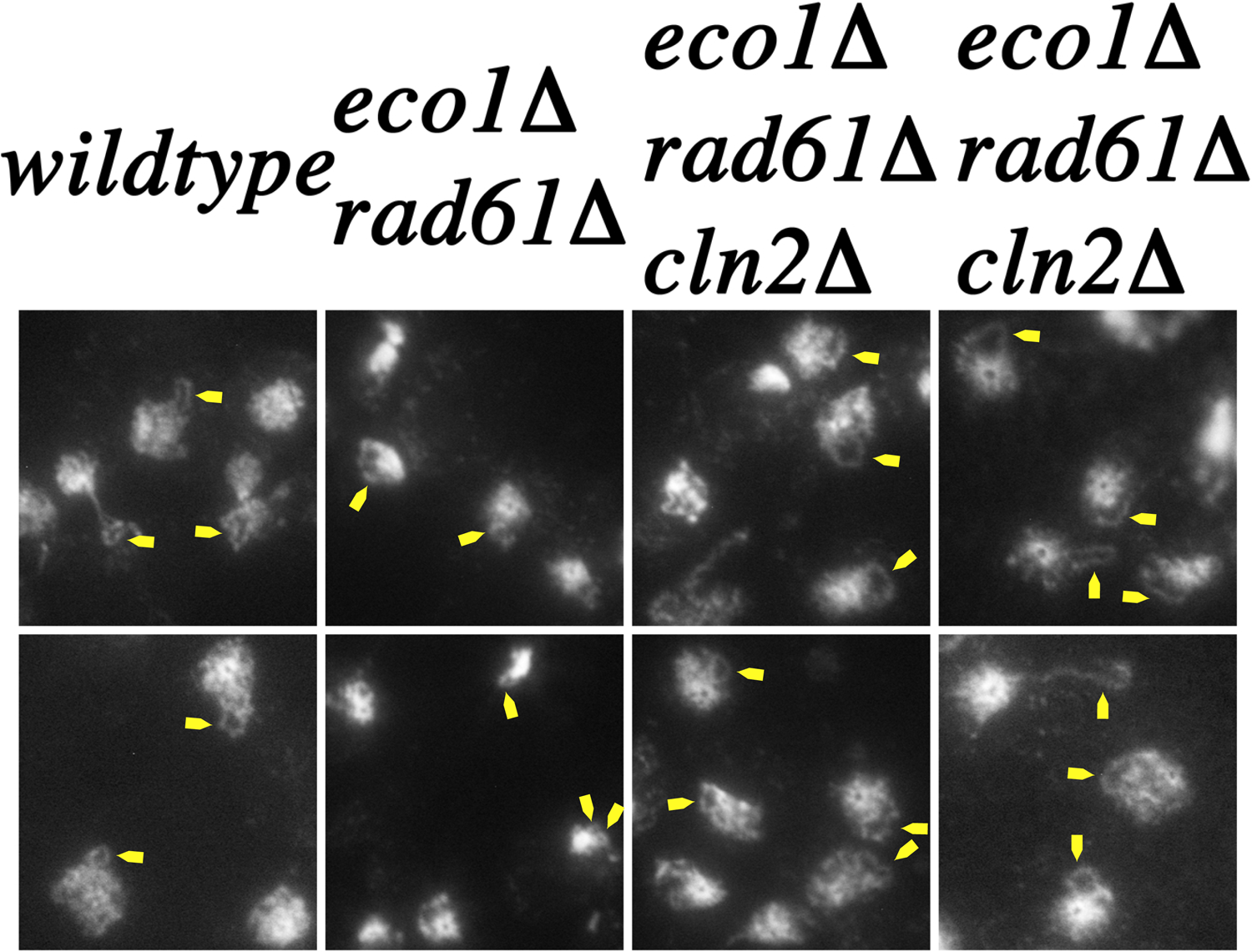
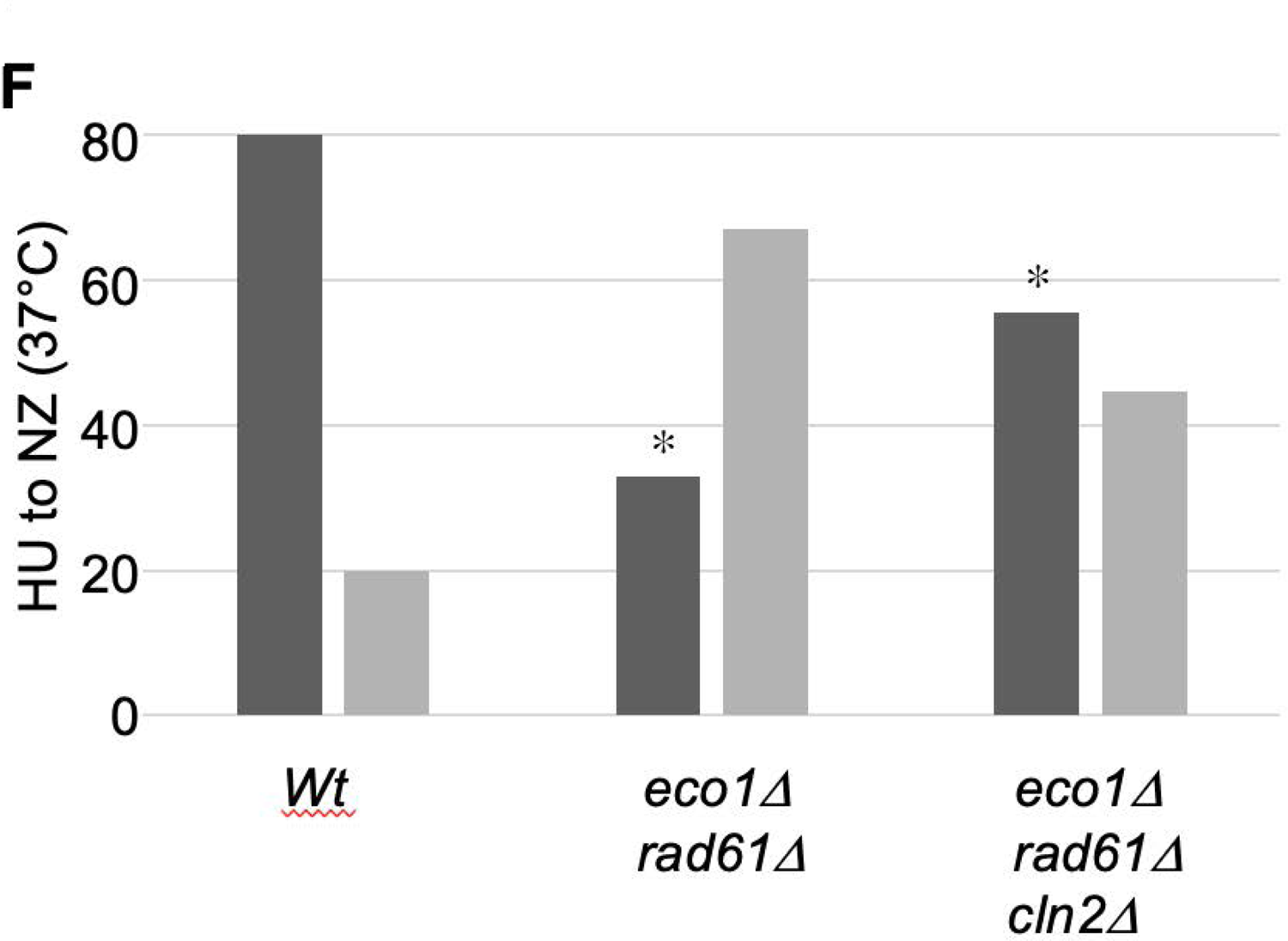
Deletion of *CLN2* rescues the chromatin hypercondensation defect otherwise present in *eco1Δ rad61Δ* cells under thermic stress. A) and D) DNA content of wildtype cells, parental *eco1Δ rad61Δ* double mutant cells, and 2 independent isolates of *eco1Δ rad61Δ cln2Δ* triple mutant at either log phase (A) or synchronized in S phase using hydroxyurea (D), prior to growth for 3 hours at 37°C in medium supplemented with nocodazole (NZ). B) and E) Micrographs of DNA masses and rDNA obtained from preanaphase cells as described in (A) and (D), respectively. Chromosomal masses and rDNA loop structures were detected using DAPI. White arrows indicate rDNA loops, which typically reside proximal to the genomic mass. C) and F) Quantification of genome mass areas, relative to template, for cells described in (A) and (D), respectively. For graph C: Wildtype (n=38), *eco1Δ rad61Δ* (n=105), *eco1Δ rad61Δ cln2Δ* (two replicates: n=237 and n=176 to produce an n_total_=413). For graph F: Wildtype (n=117), *eco1Δ rad61Δ* (n=189), *eco1Δ rad61Δ cln2Δ* (two replicates: n=381 and n=83 to produce an n_total_=464). Chi-Squared tests were subsequently used to assess the dependence of gene mutations (* indicates p-value at or below 0.05) on chromatin condensation levels. For graph C: wildtype, p-value = 0.999; *eco1Δ rad61Δ*, p-value = 0.0000272; *eco1Δ rad61Δ cln2Δ*, p-value = 0.00104. For graph F: wildtype, p-value = 0.983; *eco1Δ rad61Δ*, p-value = 3.74×10^-14^; *eco1Δ rad61Δ cln2Δ*, p-value = 4.45×10^-14^. Not shown: we also compared chromatin condensation effects across the three strains. The percentage of genomic masses, within a given field, that matched or exceeded the template was calculated and those values used to determine statistical significance: wildtype to *eco1Δ rad61Δ, p=0.00068; eco1Δ rad61Δ* to *eco1Δ rad61Δ cln2Δ* (both replicates combined), p=0.0066.

We thus focused on quantifying genomic mass areas for each of the four strains to assess the impact of *CLN2* deletion on chromosome condensation. The area of each genomic mass (excluding rDNA loops) was individually queried for all cells in the field of view, and compared to a standard area typical of DNA masses in wildtype cells (see Materials and Methods). Indeed, the results show that the majority (89%) of DNA masses from wildtype cells matched the template implemented for this analysis (Figures 6B,C). In contrast, the genomic mass area of *eco1Δ rad61Δ* double mutant cells appeared reduced and thus more highly condensed (Figure 6B), with only 30% of DNA masses matching that of the template (Figure 6C). The majority of genomic masses from *eco1Δ rad61Δ cln2*Δ triple mutant cells returned to a near wildtype level of condensation (Figure 6B) with 66% matching that of the template (Figure 6C). While the increase of genomic mass hypercondensation in *eco1Δ rad61Δ* double mutant cell was statistically significant, compared to wildtype cells, the level of genomic mass hypercondensation of *eco1Δ rad61Δ cln2Δ* triple mutant cells was also significantly different from that of wildtype cells. These results suggest that rescue of hypercondensation defects is not the only mechanism through which *CLN2* deletion promotes cell viability. Toward this end, we note that double rDNA rings, indicative of cohesion defects (Shen et al., 2017; Guacci et al., 1997), were seldom observed in *eco1Δ rad61Δ cln2Δ* triple mutant cells.

Our cell cycle experiments suggest that the positive effect of *CLN2* deletion occurs independent of the role for Cln2-CDK in START (Figure 5). If true, then triple mutant cells shifted to the restrictive temperature of 37°C after START should result in normal condensation of the genomic masses. To test this prediction, log phase cultures of wildtype cells, *eco1Δ rad61Δ* double mutant cells, and two independent isolates of *eco1Δ rad61Δ cln2Δ* triple mutant cells, were incubated at 30°C in fresh medium that contained hydroxyurea (HU), which arrests cells after START in early S phase. The resulting S phase cultures were washed, released into fresh medium supplemented with nocodazole, and incubated at 37°C for 3 hours. Cell samples were harvested at each synchronization step and DNA contents analyzed by flow cytometry (Figure 6D). As in the prior experiment, wildtype cells contained large and uniformly-stained genomic DNA masses (Figure 6E) in which the majority (80%) matched the template area (Figure 6F). Discrete loops of rDNA were also easily discernible (Figure 6E). The genomic mass areas obtained from *eco1Δ rad61Δ* double mutant cells, however, appeared more condensed (Figure 6E), such that only 33% of the DNA masses matched the template (Figure 6F). When distinguishable, the rDNA loops appeared shorter, compared to those obtained from wildtype cells (Figure 6E). Despite shifting to 37°C after START, genomic masses of *eco1Δ rad61Δ cln2Δ* triple mutant cells exhibited increased areas (Figure 6E), with 56% matching the template and thus reverting to near wildtype (Figure 6F). In addition to the rescue in genomic mass area, the rDNA appeared similar to those in wildtype cells (Figure 6E). These results suggest both that Cln2-CDK antagonizes a chromatin condensation activity in cells under hyperthermic stress and that this activity occurs independent of START regulation.

## DISCUSSION

Eco1/ESCO2 and cohesins are required for an impressive array of cellular processes: cohesion establishment coupled to DNA replication during S phase, chromosome condensation and DNA repair during G2/M phase, nucleolar integrity, chromosome orientation within the nucleus, and dynamic transcriptional regulation during G1 so that cells can respond appropriately to changing internal and external cues. The identity of Cln2 as a key regulator of cohesin-based functions represents an important step forward in deciphering chromatin biology. Beyond Cln2, only Rad61 appears to directly antagonize cohesin function (Rolef Ben Shahar et al., 2009; Sutani et al., 2009). The antagonistic roles played by Rad61/WAPL include dissociating cohesin from DNA during S phase and altering DNA extrusion loop lengths during G1 (Guacci and Koshland, 2012; Tedeschi et al., 2013; Lopez-Serra et al., 2013; Wutz et al., 2017; Haarhuis et al., 2017; Gassler et al., 2017; Bloom et al., 2018). Other gene mutations that suppress phenotypes associated with *ECO1* mutation occur in a more predictable fashion. For instance, *smc3^K113N^* is an acetyl-mimic allele, providing a straight-forward rationale for the suppression of defects exhibited by *eco1* mutant cells (Rolef Ben-Shahar et al., 2009; Sutani et al., 2009). Some *eco1* mutant cell phenotypes are suppressed by specific mutations within *PDS5*, but Pds5 serves as a bridge between Rad61. These *PDS5* mutations, however, appear to have little effect on cell growth or cohesion (Sutani et al., 2009). An intragenic allele of *ECO1* also suppresses *eco1*-dependent growth defects, likely through promoting Eco1 dimerization (Sutani et al., 2009). Note that several studies now point to the importance of both Eco1 and cohesin dimerization/clustering during the establishment of sister chromatid cohesion and DNA loop formation, although other studies suggest that at least cohesins may act in monomeric form (Cattoglio et al., 2019; Shi et al., 2020; Kulemzina et al., 2012, Tong and Skibbens, 2014; Eng et al., 2015; Zhang et al., 2008; Xiang and Koshland, 2021; Skibbens et al., 1999; Kim et al., 2019; Liu et al., 2020; Shi et al., 2020). As a final class of suppressors, *ELG1* deletion and PCNA overexpression, both of which result in elevated levels of chromatin-bound PCNA, suppress *eco1* mutant cell phenotypes through increased Eco1 recruitment to the DNA replication fork (Skibbens et al., 1999; Moldovan et al., 2006; Maradeo and Skibbens, 2010; Parnas et al., 2010; Song et al., 2012; Bender et al., 2020; Zuilkoski and Skibbens, 2020a). Cln2 appears distinct from each of these suppressor pathways and thus creates a new category of cohesin regulators.

A second major revelation of this study is the unique role, among G1 cyclins, that Cln2 exhibits on cohesin pathways. G1 cyclins are critical for the irreversible transition from G1 into S phase, or START. Cln1-Cln3 are functionally redundant such that 1) any single G1 cyclin can promote START and 2) *CLN1-CLN3* triple mutants are rescued by the expression of any one of several human G1 cyclins (Richardson et al., 1989; Lew et al., 1991; Koff et al., 1999; Lew et al., 1992; Levine et al., 1996; Queralt and Igual, 2004). G1 cyclins, however, can exert temporal differences in START regulation. For instance, Cln3 in association with cyclin-dependent kinase (Cln3-CDK) is an early inactivator of Whi5 (reviewed Fisher, 2016; Ewald, 2018; Li et al., 2021). Whi5, the yeast homolog of human retinoblastoma protein (Crane et al., 2019), is an inhibitor of the SBF-dependent transcription of genes that are required for the transition into S phase (Talerak et al., 2017; Teufel et al., 2019). SBF (and MBF) activation results in Cln1 and Cln2 upregulation, leading to Cln1-CDK and Cln2-CDK complexes that further inactivate Whi5 and provide a positive feedback mechanism to promote START. Due to the asymmetric division that occurs in *Saccharomyces cerevisiae*, the additional growth phase required by small buds (as opposed to the larger mother cells) that result from cell division appears more dependent on the shared activities of Cln1-CDK and Cln2-CDK. Moreover, Cln1 and Cln2 proteins are roughly 60% identical at the amino acid level and exhibit nearly identical target docking-site motifs (Bhaduri et al., 2015; Bandyopadhyay et al., 2020). Little if any evidence differentiates the activities of Cln1-CDK and Cln2-CDK. Thus, our findings that deletion of *CLN2*, but not that of either *CLN1* or *CLN3*, suppresses the growth defect of *eco1Δ rad61Δ* double mutant cells highlight an important distinction between G1-CDKs and reveal a previously unknown role for Cln2-CDK. The identification of Cln2-CDK substrates that render *eco1Δ rad61Δ* double mutant cells inviable under thermic stress remains unknown but likely involves cohesin modifications given that both Eco1 and Rad61 regulate cohesins. In contrast, heat shock/chaperone proteins Hsp80 and Hsc82 also regulate hyperthermic-induced rDNA hyper-condensation, although this condensed chromatin state does not overtly extend to the remaining genomic mass (Shen and Skibbens, 2020). Condensin complexes (condensin I and II in vertebrate cells) represent another set of factors through which chromosome condensation is regulated during G2/M and thus could be targets of Cln2-CDK (Davidson et al., 2021; Skibbens 2019; Paulson et al., 2021). Testing which of these factors (cohesins, HSPs, condensins, or other complex components) are targeted by Cln2-CDK during hyperthermic stress (as revealed in *eco1Δ rad61Δ* mutant cells) represents a significant undertaking for future studies, but the findings presented here regarding the dramatic changes in chromatin structure provide clues critical for directing those efforts.

The revelation that *CLN2* deletion fully rescues *eco1 rad61* mutant cell ts growth defects (and suppresses *eco1* mutant ts growth defects), is somewhat surprising. For instance, *eco1* mutation renders cells inviable when combined with mutated alleles of either *CAK1* (encoding the CDK-activating kinase Cak1) or *CDC28* (encoding the catalytic subunit of all Cyclin-Dependent Kinases in budding yeast) (Brands and Skibbens, 2008). Results from that study further revealed that CDKs indeed promote cohesion. *A priori*, these findings predict that cyclin mutation (or deletion) would be lethal in combination with *eco1* mutations, in contrast to the rescue reported here. Resolving this apparent conundrum likely involves differentiating between the roles of mitotic CDK (deficiencies which are lethal in *eco1* mutant cells) from Cln2-CDK (deficiencies which suppress *eco1* mutant cell growth defects). The intersection of Eco1 and CDKs, however, is even more complex (Figure 7). Elegant studies revealed that Eco1 is a phosphoprotein and CDK substrate (Ubersax et al., 2003), which is part of a signal that promotes Eco1 degradation during G2 and limits cohesion establishment activities to S phase (Skibbens et al., 1999; Lyons and Morgan, 2011; Lyons et al., 2013; Seoane and Morgan, 2017). This CDK/phosphorylation-dependent mechanism appears highly conserved, with subsequent ubiquitination provided by the Skp1–Cullin– F-box (SCF) ubiquitin ligase and augmented by Cullin Ring-like (CRL4) ubiquitin ligase (Lyons et al., 2013; Minamino et al. 2018; Sun et al., 2019). The mechanism through which *CLN2* deletion suppresses *eco1Δ rad61Δ* mutant cell ts phenotypes, however, is demonstrably distinct from the CDK-dependent mechanism through which Eco1 activity is limited - given that *ECO1* is fully deleted from these cells.

**Figure 7.**
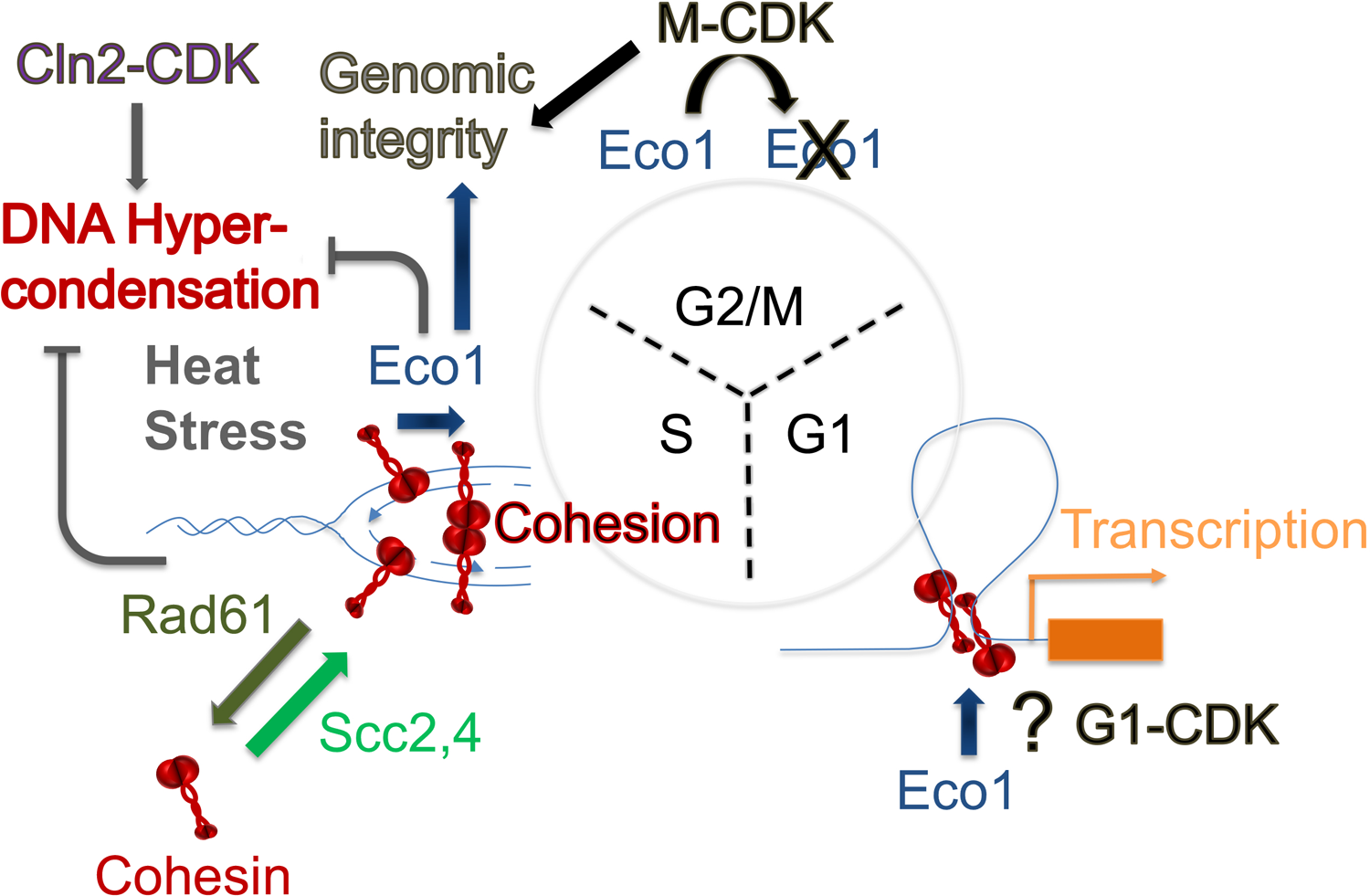
Multi-faceted and opposing roles for CDK regulation of Eco1 activity. In this current study, we provide evidence that Cln2-CDK regulates chromatin condensation reactions in combination with Eco1 and Rad61. Cln2-CDK promotes chromatin hyper-condensation after START (possibly during S phase) and in response to thermic stress. Also during S phase, Eco1 establishes cohesion between cohesins (in red) newly deposited (via Scc2,4) onto nascent sister chromatids after passage of the DNA replication fork. Eco1 acetylation of cohesin (stabilizing cohesin dimers, red) blocks Rad61-dependent cohesin dissociation activities. After S phase, M-CDKs phosphorylate Eco1, resulting in Eco1 degradation through both G2 and M phases. In response to damage, however, Eco1 expression is significantly upregulated during G2/M to promote DNA damage-induced cohesion (not shown), even as Eco1 degradation continues. Intriguingly, *eco1* is synthetically lethal in combination with either *cdc28* (CDK) or *cak* (CDK activating kinase), revealing that Eco1 and M-CDK perform complementary roles to maintain genomic integrity (which are likely to include chromosome condensation, cohesion maintenance, and transcription regulation). During G1, Eco1 is critical for regulation gene transcription and genomic architecture within the nucleus. The role of G1-CDKs in this portion of the cell cycle remain largely unknown.

What is the cellular process through which *CLN2* deletion rescues *eco1Δ rad61Δ* mutant cell ts lethality? While the current study focused on chromosome hypercondensation, structural changes in chromatin are likely to also impact gene transcription. Notably, transcriptional dysregulations that arise due to mutation of either *ESCO2* or cohesin genes are typically lethal in humans (see below). We also note the relative absence of cohesion defects, as evidenced by lack of obvious rDNA double rings in *eco1Δ rad61Δ cln2Δ* triple mutant cells. Thus, our findings do not exclude the possibilities that Cln2, through CDK activation, impacts numerous cohesin functions through which cohesion, condensation, transcription and DNA repair are mediated. Our results do reveal, however, that regulating the various cohesin functions are parsed to different factors, consistent with prior observations (Guacci and Koshland, 2012; Zuilkoski and Skibbens 2020a).

Finally, the identification of *CLN2* deletion as a suppressor of *eco1 rad61* mutant cell ts growth defects provides a new framework for interpreting results from studies regarding the role of cyclins in development and cancer. In humans, severe developmental defects, collectively referred to as cohesinopathies, result from mutation of *ESCO2* (homolog of yeast Eco1/Ctf7), most cohesin genes (*SMC1A*, *SMC3*, *PDS5*, *RAD21*) or cohesin regulators (*NIPBL* or *HDAC8*) result in (Banerji et al., 2017; Sarogni et al., 2020; Mfarej and Skibbens, 2020; Vega et al., 2020). All of these cohesin-pathway genes are essential, such that the syndromes that arise from *ESCO2* mutation (Roberts Syndrome, RBS) or cohesin/regulator mutations (Cornelia de Lange Syndrome, CdLS) occur infrequently - likely due to early and spontaneous pregnancy terminations. Individuals with RBS and CdLS exhibit defects in craniofacial structures, limb growth, GI and respiratory tracts, heart formation, and even intellectual abilities (Smithellis and Newman, 1992). Of interest here is that *SMC1A* mutated CdLS patient cells often exhibit down-regulation of a G1 cyclin (cyclin D1, Ccnd1). While this observation prompted those authors to link reduced Ccdn1 levels to apoptosis, cell cycle delay, and tumorigenesis (Fazio et al., 2016), our findings formally raise the possibility that CdLS patient cell survival may result in part from Ccnd1 reduction, a model supported by findings that *nipblb/smc1a* reduction does not necessarily produce Ccnd1 down regulation (Fazio et al., 2016). On the other hand, the upregulation of Ccnd1 is tightly correlated with melanoma, breast, and other cancers (Gonzalez-Ruiz et al., 2021; Ramos-Garcia et al., 2019). Future studies focused on the intersection between human Cln2 homologs, Esco2/cohesin pathways, and changes in chromatin structure will likely provide new insights into both tumorigenesis and birth defects.

## ACKNOWLEDGEMENTS

We thank Dr. Meg Kenna and the many Skibbens lab members (Drs. Michael Mfarej, Caitlyn Zuilkoski, Donglai Shen, Raj Benerji, Kevin Tong, Soumya Rudra, Marie Maradeo, Christina Sie, and also Annie Sanchez, Gurvir Singh, Nicole Kirven, Caitlyn Devine, Ariana Malik, Emma Anderson, Chris Geissler, Shaya Ameri, Samantha Sarli, Rachel Sternberg and Divya Sirdeshpandi, Krupa Patel and Anne Smolko) that were present over the course of this work and contributed their thoughts and support. We are deeply indebted to Drs. Gregory Lang and Vincent Guacci for the sharing of reagents and expertise.

## REFERENCES

1. Adane B, Alexe G, Seong BKA, Lu D, Hwang EE, Hnisz D, Lareau CA, Ross L, Lin S, Dela Cruz FS, Richardson M, Weintraub AS, Wang S, Iniguez AB, Dharia NV, Conway AS, Robichaud AL, Tanenbaum B, Krill-Burger JM, Vazquez F, Schenone M, Berman JN, Kung AL, Carr SA, Aryee MJ, Young RA, Crompton BD, and Stegmaier K. 2021. STAG2 loss rewires oncogenic and developmental programs to promote metastasis in Ewing sarcoma. Cancer Cell. 39(6):827–844.

2. Alomer RM, da Silva EML, Chen J, Piekarz KM, McDonald K, Sansam CG, Sansam CL, and Rankin S. 2017. Esco1 and Esco2 regulate distinct cohesin functions during cell cycle progression. Proc Natl Acad Sci U S A. 114(37):9906–9911.

3. Bandyopadhyay S, Bhaduri S, Örd M, Davey NE, Loog M, and Pryciak PM. 2020. Comprehensive Analysis of G1 Cyclin Docking Motif Sequences that Control CDK Regulatory Potency In Vivo. Curr Biol. 30(22):4454–4466.

4. Banerji R, Eble DM, Iovine MK, and Skibbens RV. 2016. Esco2 regulates cx43 expression during skeletal regeneration in the zebrafish fin. Dev Dyn. 245(1):7–21.

5. Banerji R, Skibbens RV, and Iovine MK. 2017. How many roads lead to cohesinopathies? Dev Dyn. 246(11):881–888.

6. Banerji R, Skibbens RV, and Iovine MK. 2017. Cohesin mediates Esco2-dependent transcriptional regulation in a zebrafish regenerating fin model of Roberts Syndrome. Biol Open. 6(12):1802–1813.

7. Bellows AM, Kenna MA, Cassimeris L, and Skibbens RV. 2003. Human EFO1p exhibits acetyltransferase activity and is a unique combination of linker histone and Ctf7p/Eco1p chromatid cohesion establishment domains. Nucleic Acids Res. 31(21):6334–6343.

8. Bender D, Da Silva EML, Chen J, Poss A, Gawey L, Rulon Z, and Rankin S. 2020. Multivalent interaction of ESCO2 with the replication machinery is required for sister chromatid cohesion in vertebrates. Proc Natl Acad Sci U S A. 117(2):1081–1089.

9. Bhaduri S, Valk E, Winters MJ, Gruessner B, Loog M, and Pryciak PM. 2015. A docking interface in the cyclin Cln2 promotes multi-site phosphorylation of substrates and timely cell-cycle entry. Curr Biol. 25(3):316–325.

10. Billon P, Li J, Lambert JP, Chen Y, Tremblay V, Brunzelle JS, Gingras AC, Verreault A, Sugiyama T, Couture JF, and Côté J. 2017. Acetylation of PCNA Sliding Surface by Eco1 Promotes Genome Stability through Homologous Recombination. Mol Cell. 65(1):78–90.

11. Bloom MS, Koshland D, and Guacci V. 2018. Cohesin Function in Cohesion, Condensation, and DNA Repair Is Regulated by Wpl1p via a Common Mechanism in Saccharomyces cerevisiae. Genetics. 208(1):111-124.

12. Boginya A, Detroja R, Matityahu A, Frenkel-Morgenstern M, and Onn I. 2019. The chromatin remodeler Chd1 regulates cohesin in budding yeast and humans. Sci Rep. 9(1):8929.

13. Brands A, and Skibbens RV. 2008. Sister chromatid cohesion role for CDC28-CDK in Saccharomyces cerevisiae. Genetics. 180(1):7–16.

14. Chin CV, Antony J, Ketharnathan S, Labudina A, Gimenez G, Parsons KM, He J, George AJ, Pallotta MM, Musio A, Braithwaite A, Guilford P, Hannan RD, and Horsfield JA. 2020. Cohesin mutations are synthetic lethal with stimulation of WNT signaling. Elife. 9:e61405.

15. Cingolani P, Platts A, Wang LL, Coon M, Nguyen T, Wang L, Land SJ, Lu X, and Ruden DM. 2012. A program for annotating and predicting the effects of single nucleotide polymorphisms, SnpEff: SNPs in the genome of Drosophila melanogaster strain w1118; iso-2; iso-3.”, Fly (Austin) 6(2):80-92.

16. Ciosk R, Shirayama M, Shevchenko A, Tanaka T, Toth A, Shevchenko A, and Nasmyth K. 2000. Cohesin’s binding to chromosomes depends on a separate complex consisting of Scc2 and Scc4 proteins. Mol Cell. 5(2):243–254.

17. Covo S, Westmoreland JW, Gordenin DA, and Resnick MA. 2010. Cohesin Is limiting for the suppression of DNA damage-induced recombination between homologous chromosomes. PLoS Genet. 6(7):e1001006.

18. Crane MM, Tsuchiya M, Blue BW, Almazan JD, Chen KL, Duffy SR, Golubeva A, Grimm AM, Guard AM, Hill SA, Huynh E, Kelly RM, Kiflezghi M, Kim HD, Lee M, Lee TI, Li J, Nguyen BMG, Whalen RM, Yeh FY, McCormick M, Kennedy BK, Delaney JR, and Kaeberlein M. 2019. Rb analog Whi5 regulates G1 to S transition and cell size but not replicative lifespan in budding yeast. Transl Med Aging. 3:104–108.

19. D’Ambrosio C, Kelly G, Shirahige K, and Uhlmann F. 2008. Condensin-dependent rDNA decatenation introduces a temporal pattern to chromosome segregation. Curr Biol. 18(14):1084–1089.

20. Davidson MB, Katou Y, Keszthelyi A, Sing TL, Xia T, Ou J, Vaisica JA, Thevakumaran N, Marjavaara L, Myers CL, Chabes A, Shirahige K, and Brown GW. 2012. Endogenous DNA replication stress results in expansion of dNTP pools and a mutator phenotype. EMBO J. 31(4):895–907.

21. Davidson IF, Goetz D, Zaczek MP, Molodtsov MI, Huis In’t Veld PJ, Weissmann F, Litos G, Cisneros DA, Ocampo-Hafalla M, Ladurner R, Uhlmann F, Vaziri A, and Peters JM. 2016. Rapid movement and transcriptional re-localization of human cohesin on DNA. EMBO J. 35(24):2671-2685.

22. Davidson IF, and Peters JM. 2021. Genome folding through loop extrusion by SMC complexes. Nat Rev Mol Cell Biol. 22(7):445–464.

23. de Los Santos-Velázquez AI, de Oya IG, Manzano-López J, and Monje-Casas F. 2017. Late rDNA Condensation Ensures Timely Cdc14 Release and Coordination of Mitotic Exit Signaling with Nucleolar Segregation. Curr Biol 27(21):3248–3263.

24. de Wit E, Vos ES, Holwerda SJ, Valdes-Quezada C, Verstegen MJ, Teunissen H, Splinter E, Wijchers PJ, Krijger PH, and de Laat W. 2015. CTCF Binding Polarity Determines Chromatin Looping. Mol Cell. 60(4):676–684.

25. Deardorff MA, Noon SE, and Krantz ID. 2020. Cornelia de Lange Syndrome. In: Adam MP, Ardinger HH, Pagon RA, Wallace SE, Bean LJH, Gripp KW, Mirzaa GM, Amemiya A, editors. GeneReviews® [Internet]. Seattle (WA): University of Washington, Seattle;1993–2022.

26. Ding DQ, Sakurai N, Katou Y, Itoh T, Shirahige K, Haraguchi T, and Hiraoka Y. 2006. Meiotic cohesins modulate chromosome compaction during meiotic prophase in fission yeast. J Cell Biol. 174(4):499–508.

27. Dorsett D. 2016. The Drosophila melanogaster model for Cornelia de Lange syndrome: Implications for etiology and therapeutics. Am J Med Genet C Semin Med Genet. 172(2):129–137.

28. Eng T, Guacci V, and Koshland D. 2015. Interallelic complementation provides functional evidence for cohesin-cohesin interactions on DNA. Mol Biol Cell. 26(23):4224–4235.

29. Engel SR, Dietrich FS, Fisk DG, Binkley G, Balakrishnan R, Costanzo MC, Dwight SS, Hitz BC, Karra K, Nash RS, Weng S, Wong ED, Lloyd P, Skrzypek MS, Miyasato SR, Simison M, and Cherry JM. 2013. The Reference Genome Sequence of Saccharomyces cerevisiae: Then and Now. G3 (Bethesda). pii: g3.113.008995v1. doi: 10.1534/g3.113.008995. PMID:24374639.

30. Ewald JC. 2018. How yeast coordinates metabolism, growth and division. Curr Opin Microbiol. 45:1–7.

31. Fazio G, Gaston-Massuet C, Bettini LR, Graziola F, Scagliotti V, Cereda A, Ferrari L, Mazzola M, Cazzaniga G, Giordano A, Cotelli F, Bellipanni G, Biondi A, Selicorni A, Pistocchi A, and Massa V. 2016. CyclinD1 Down-Regulation and Increased Apoptosis Are Common Features of Cohesinopathies. J Cell Physiol. 231(3):613–622.

32. Fisher RP. 2016. Getting to S: CDK functions and targets on the path to cell-cycle commitment. F1000Res. 26(5):2374.

33. Gandhi R, Gillespie PJ, and Hirano T. 2006. Human Wapl is a cohesin-binding protein that promotes sister-chromatid resolution in mitotic prophase. Curr Biol. 6(24):2406–2417.

34. Gard S, Light W, Xiong B, Bose T, McNairn AJ, Harris B, Fleharty B, Seidel C, Brickner JH, and Gerton JL. 2009. Cohesinopathy mutations disrupt the subnuclear organization of chromatin. J Cell Biol. 187(4):455–462.

35. Garrison E, and Marth G. 2012. Haplotype-based variant detection from short-read sequencing. arXiv preprint arXiv:1207.3907 [q-bio.GN].

36. Gassler J, Brandão HB, Imakaev M, Flyamer IM, Ladstätter S, Bickmore WA, Peters JM, Mirny LA, and Tachibana K. 2017. A mechanism of cohesin-dependent loop extrusion organizes zygotic genome architecture. EMBO J. 36(24):3600–3618.

37. Gentekaki E, Curtis BA, Stairs CW, Klimeš V, Eliáš M, Salas-Leiva DE, Herman EK, Eme L, Arias MC, Henrissat B, Hilliou F, Klute MJ, Suga H, Malik SB, Pightling AW, Kolisko M, Rachubinski RA, Schlacht A, Soanes DM, Tsaousis AD, Archibald JM, Ball SG, Dacks JB, Clark CG, van der Giezen M, and Roger AJ. 2017. Extreme genome diversity in the hyper-prevalent parasitic eukaryote Blastocystis. PLoS Biol. 15(9):e2003769.

38. González-Ruiz L, González-Moles MÁ, González-Ruiz I, Ruiz-Ávila I, and Ramos-García P. 2021. Prognostic and Clinicopathological Significance of CCND1/Cyclin D1 Upregulation in Melanomas: A Systematic Review and Comprehensive Meta-Analysis Cancers (Basel) 13(6):1314.

39. Gordillo M, Vega H, Trainer AH, Hou F, Sakai N, Luque R, Kayserili H, Basaran S, Skovby F, Hennekam RC, Uzielli ML, Schnur RE, Manouvrier S, Chang S, Blair E, Hurst JA, Forzano F, Meins M, Simola KO, Raas-Rothschild A, Schultz RA, McDaniel LD, Ozono K, Inui K, Zou H, and Jabs EW. 2008. The molecular mechanism underlying Roberts syndrome involves loss of ESCO2 acetyltransferase activity. Hum Mol Genet. 17(14):2172–80.

40. Guacci V, Hogan E, and Koshland D. 1994. Chromosome condensation and sister chromatid pairing in budding yeast. J Cell Biol. 125(3):517–530.

41. Guacci V, Koshland D, and Strunnikov A. 1997. A direct link between sister chromatid cohesion and chromosome condensation revealed through the analysis of MCD1 in S. cerevisiae. Cell. 91(1):47–57.

42. Guacci V, and Koshland D. 2012. Cohesin-independent segregation of sister chromatids in budding yeast. Mol Biol Cell. 23(4):729–739.

43. Haarhuis JHI, van der Weide RH, Blomen VA, Yáñez-Cuna JO, Amendola M, van Ruiten MS, Krijger PHL, Teunissen H, Medema RH, van Steensel B, Brummelkamp TR, de Wit E, and Rowland BD. 2017. The Cohesin Release Factor WAPL Restricts Chromatin Loop Extension. Cell. 169(4):693–707.

44. Heidinger-Pauli JM, Unal E, Guacci V, and Koshland D. 2008. The kleisin subunit of cohesin dictates damage-induced cohesion. Mol Cell. 31(1):47–56.

45. Heidinger-Pauli JM, Unal E, and Koshland D. 2009. Distinct targets of the Eco1 acetyltransferase modulate cohesion in S phase and in response to DNA damage. Mol Cell. 34(3):311–321.

46. Horsfield JA, Anagnostou SH, Hu JK, Cho KH, Geisler R, Lieschke G, Crosier KE, and Crosier PS. 2007. Cohesin-dependent regulation of Runx genes. Development. 134(14):2639–2649.

47. Horsfield JA. 2022. Full circle: a brief history of cohesin and the regulation of gene expression. FEBS J. doi:10.1111/febs.16362.

48. Ito T, Ando H, Suzuki T, Ogura T, Hotta K, Imamura Y, Yamaguchi Y, and Handa H. 2010. Identification of a primary target of thalidomide teratogenicity. Science. 327:1345–1350.

49. Ivanov D, Schleiffer A, Eisenhaber F, Mechtler K, Haering CH, and Nasmyth K. 2002. Eco1 is a novel acetyltransferase that can acetylate proteins involved in cohesion. Curr Biol. 12(4):323–328.

50. Johnson C, Gali VK, Takahashi TS, and Kubota T. 2016. PCNA Retention on DNA into G2/M Phase Causes Genome Instability in Cells Lacking Elg1. Cell Rep. 16(3):684–695.

51. Kakui Y, and Uhlmann F. 2018. SMC complexes orchestrate the mitotic chromatin interaction landscape. Curr Genet. 64(2):335–339.

52. Kawauchi S, Calof AL, Santos R, Lopez-Burks ME, Young CM, Hoang MP, Chua A, Lao T, Lechner MS, Daniel JA, Nussenzweig A, Kitzes L, Yokomori K, Hallgrimsson B, and Lander AD. 2009. Multiple organ system defects and transcriptional dysregulation in the Nipbl(+/-) mouse, a model of Cornelia de Lange Syndrome. PLoS Genet. 5(9):e1000650.

53. Ketharnathan S, Labudina A, and Horsfield JA. 2020. Cohesin Components Stag1 and Stag2 Differentially Influence Haematopoietic Mesoderm Development in Zebrafish Embryos. Front Cell Dev Biol. 8:617545.

54. Kim JS, Krasieva TB, LaMorte V, Taylor AM, and Yokomori KJ. 2002. Specific recruitment of human cohesin to laser-induced DNA damage. Biol Chem. 277(47):45149–45153.

55. Kim Y, Shi Z, Zhang H, Finkelstein IJ, and Yu H. 2019. Human cohesin compacts DNA by loop extrusion. Science. 366(6471):1345–1349.

56. Koff A, Cross F, Fisher A, Schumacher J, Leguellec K, Philippe M, and Roberts JM. 1999. Human cyclin E, a new cyclin that interacts with two members of the CDC2 gene family. Cell. 66(6):1217–1228.

57. Kong X, Ball Jr AR, Pham HX, Zeng W, Chen HY, Schmiesing JA, Kim JS, Berns M, and Yokomori K. 2014. Distinct functions of human cohesin-SA1 and cohesin-SA2 in double-strand break repair. Mol Cell Biol. 34(4):685–698.

58. Krantz ID, McCallum J, DeScipio C, Kaur M, Gillis LA, Yaeger D, Jukofsky L, Wasserman N, Bottani A, Morris CA, Nowaczyk MJ, Toriello H, Bamshad MJ, Carey JC, Rappaport E, Kawauchi S, Lander AD, Calof AL, Li HH, Devoto M, and Jackson LG. 2004. Cornelia de Lange syndrome is caused by mutations in NIPBL, the human homolog of Drosophila melanogaster Nipped-B. Nat Genet. 36(6):631–635.

59. Kueng S, Hegemann B, Peters BH, Lipp JJ, Schleiffer A, Mechtler K, and Peters JM. 2006. Wapl controls the dynamic association of cohesin with chromatin. Cell. 127(5):955–967.

60. Kulemzina I, Schumacher MR, Verma V, Reiter J, Metzler J, Failla AV, Lanz C, Sreedharan VT, Rätsch G, and Ivanov D. 2012. Cohesin rings devoid of Scc3 and Pds5 maintain their stable association with the DNA. PLoS Genet. 8(8):e1002856.

61. Lamothe R, Costantino L, and Koshland DE. 2020. The spatial regulation of condensin activity in chromosome condensation. Genes Dev. 34(11-12):819–831.

62. Lavoie BD, Hogan E, and Koshland D. 2004. In vivo requirements for rDNA chromosome condensation reveal two cell-cycle-regulated pathways for mitotic chromosome folding. Genes Dev. 18(1):76–87.

63. Levine K, Huang K, and Cross FR. 1996. Saccharomyces cerevisiae G1 cyclins differ in their intrinsic functional specificities. Mol Cell Biol. 16(12):6794–6803.

64. Lew DJ, Dulić V, and Reed SI. 1991. Isolation of three novel human cyclins by rescue of G1 cyclin (Cln) function in yeast. Cell. 66(6):1197–1206.

65. Lew DJ, Marini NJ, and Reed SI. 1992. Different G1 cyclins control the timing of cell cycle commitment in mother and daughter cells of the budding yeast S. cerevisiae. Cell. 69(2):317–327.

66. Li H, and Durbin R. 2010. Fast and accurate long-read alignment with Burrows-Wheeler Transform. Bioinformatics, Epub. [PMID: 20080505]

67. Li P, Zhimin H, and Zeng F. 2021. Tumor suppressor stars in yeast G1/S transition. Curr Genet. 67(2):207–212.

68. Lightfoot J, Testori S, Barroso C, and Martinez-Perez E. 2011. Loading of meiotic cohesin by SCC-2 is required for early processing of DSBs and for the DNA damage checkpoint. Curr Biol. 21(17):1421–1430.

69. Liu W, Biton E, Pathania A, Matityahu A, Irudayaraj J, and Onn I. 2020. Monomeric cohesin state revealed by live-cell single-molecule spectroscopy. EMBO Rep. 21(2):e48211.

70. Longtine MS, McKenzie A 3rd, Demarini DJ, Shah NG, Wach A, Brachat A, Philippsen P, and Pringle JR. 1998. Additional modules for versatile and economical PCR-based gene deletion and modification in Saccharomyces cerevisiae. Yeast. 14(10):953-961.

71. Lopez-Serra L, Lengronne A, Borges V, Kelly G, and Uhlmann F. 2013. Budding yeast Wapl controls sister chromatid cohesion maintenance and chromosome condensation. Curr Biol. 23(1):64–69.

72. Lu S, Goering M, Gard S, Xiong B, McNairn AJ, Jaspersen SL, and Gerton JL. 2010. Eco1 is important for DNA damage repair in S. cerevisiae. Cell Cycle. 9(16):3315–3327.

73. Lyons NA, and Morgan DO. 2011. Cdk1-dependent destruction of Eco1 prevents cohesion establishment after S phase. Mol Cell. 42(3):378–389.

74. Lyons NA, Fonslow BR, Diedrich JK, Yates JR 3rd, and Morgan DO. 2013. Sequential primed kinases create a damage-responsive phosphodegron on Eco1. Nat Struct Mol Biol. 20(2):194-201.

75. Maradeo ME, and Skibbens RV. 2010. Replication factor C complexes play unique pro- and anti-establishment roles in sister chromatid cohesion. PLoS One. 5(10):e15381.

76. Marsman J, O’Neill AC, Kao BR, Rhodes JM, Meier M, Antony J, Mönnich M, and Horsfield JA. 2014. Cohesin and CTCF differentially regulate spatiotemporal runx1 expression during zebrafish development. Biochim Biophys Acta. 1839(1):50–61.

77. Matos-Perdomo E, and Machín F. 2018. The ribosomal DNA metaphase loop of Saccharomyces cerevisiae gets condensed upon heat stress in a Cdc14-independent TORC1-dependent manner. Cell Cycle. 17:200–215.

78. Mfarej MG, and Skibbens RV. 2020. DNA damage induces Yap5-dependent transcription of ECO1/CTF7 in Saccharomyces cerevisiae. PLoS One. 15(12):e0242968.

79. Mfarej MG, and Skibbens RV. 2020. An ever-changing landscape in Roberts syndrome biology: Implications for macromolecular damage. PLoS Genet. 16(12):e1009219.

80. Mfarej MG, and Skibbens RV. 2022. Genetically induced redox stress occurs in a yeast model for Roberts syndrome. G3 (Bethesda). 12(2):jkab426.

81. Michaelis C, Ciosk R, and Nasmyth K. 1997. Cohesins: chromosomal proteins that prevent premature separation of sister chromatids. Cell. 91(1):35–45.

82. Minamino M, Tei S, Negishi L, Kanemaki MT, Yoshimura A, Sutani T, Bando M, and Shirahige K. 2018. Temporal Regulation of ESCO2 Degradation by the MCM Complex, the CUL4-DDB1-VPRBP Complex, and the Anaphase-Promoting Complex. Curr Biol. 28(16):2665–2672.

83. Moldovan GL, Pfander B, and Jentsch S. 2006. PCNA controls establishment of sister chromatid cohesion during S phase. Mol Cell. 23(5):723–732.

84. Mönnich M, Kuriger Z, Print CG, and Horsfield JA. 2011. A zebrafish model of Roberts syndrome reveals that Esco2 depletion interferes with development by disrupting the cell cycle. PLoS One. 6(5):e20051.

85. Parnas O, Zipin-Roitman A, Pfander B, Liefshitz B, Mazor Y, Ben-Aroya S, Jentsch S, and Kupiec M. 2010. Elg1, an alternative subunit of the RFC clamp loader, preferentially interacts with SUMOylated PCNA. EMBO J. 29(15):2611–2622.

86. Paulson JR, Hudson DF, Cisnerso-Soberanis F, and Earnshaw WC. 2021. Mitotic chromosomes. Semin Cell Dev Biol. 117:7–29.

87. Perea-Resa C, Wattendorf L, Marzouk S, and Blower MD. 2021. Cohesin: behind dynamic genome topology and gene expression reprogramming. Trends Cell Biol. 31(9):760–773.

88. Piazza A, Bordelet H, Dumont A, Thierry A, Savocco J, Girard F, and Koszul R. 2021. Cohesin regulates homology search during recombinational DNA repair. Nat Cell Biol. 23(11):1176–1186.

89. Pope PA, Bhaduri S, and Pryciak PM. 2014. Regulation of cyclin-substrate docking by a G1 arrest signaling pathway and the Cdk inhibitor Far1. Curr Biol. 24(12):1390–1396.

90. Queralt E, and Igual JC. 2004. Functional distinction between Cln1p and Cln2p cyclins in the control of the Saccharomyces cerevisiae mitotic cycle. Genetics. 168(1):129–140.

91. Ramos-García P, González-Moles MÁ, Ayén Á, González-Ruiz L, Gil-Montoya JA, and Ruiz-Ávila I. 2019. Predictive value of CCND1/cyclin D1 alterations in the malignant transformation of potentially malignant head and neck disorders: Systematic review and meta-analysis. Head Neck. 41(9):3395–3407.

92. Rao SSP, Huang SC, Glenn St Hilaire B, Engreitz JM, Perez EM, Kieffer-Kwon KR, Sanborn AL, Johnstone SE, Bascom GD, Bochkov ID, Huang X, Shamim MS, Shin J, Turner D, Ye Z, Omer AD, Robinson JT, Schlick T, Bernstein BE, Casellas R, Lander ES, Aiden EL. 2017.Cohesin Loss Eliminates All Loop Domains. Cell. 171(2):305-320.

93. Rhodes JM, Bentley FK, Print CG, Dorsett D, Misulovin Z, Dickinson EJ, Crosier KE, Crosier PS, and Horsfield JA. 2010. Positive regulation of c-Myc by cohesin is direct, and evolutionarily conserved. Dev Biol. 344(2):637–649.

94. Rogers CH, Mielczarek O, and Corcoran AE. 2021. Dynamic 3D locus organization and its drivers underpin immunoglobulin recombination. Front Immunol. 11:633705.

95. Richardson H, Wittenberg K, Cross F, and Reed S. 1989. An essential G1 function for cyclin-like proteins in yeast. Cell. 59:1127–1133.

96. Rolef Ben-Shahar T, Heeger S, Lehane C, East P, Flynn H, Skehel M, and Uhlmann F. 2008. Eco1-dependent cohesin acetylation during establishment of sister chromatid cohesion. Science. 321(5888):563–566.

97. Rollins RA, Morcillo P, and Dorsett D. 1999. Nipped-B, a Drosophila homologue of chromosomal adherins, participates in activation by remote enhancers in the cut and Ultrabithorax genes. Genetics. 152(2):577–593.

98. Rowland BD, Roig MB, Nishino T, Kurze A, Uluocak P, Mishra A, Beckouët F, Underwood P, Metson J, Imre R, Mechtler K, Katis VL, and Nasmyth K. 2009. Building sister chromatid cohesion: smc3 acetylation counteracts an antiestablishment activity. Mol Cell. 33(6):763–774.

99. Rudra S, and Skibbens RV. 2013. Cohesin codes - interpreting chromatin architecture and the many facets of cohesin function. J Cell Sci. 126(Pt 1):31–41.

100. Sanchez AC, Thren ED, Iovine MK, and Skibbens RV. 2022. Esco2 and cohesin regulate CRL4 ubiquitin ligase ddb1 expression and thalidomide teratogenicity. Cell Cycle. 6:1–13.

101. Sarogni P, Pallotta MM, and Musio A. 2020. Cornelia de Lange syndrome: from molecular diagnosis to therapeutic approach. J Med Genet. 57(5):289–295.

102. Scherzer M, Giordano F, Ferran MS, and Ström L. 2022. Recruitment of Scc2/4 to double-strand breaks depends on gammaH2A and DNA end resection. Life Sci Alliance. 5(5):e202101244.

103. Schwarzer W, Abdennur N, Goloborodko A, Pekowska A, Fudenberg G, Loe-Mie Y, Fonseca NA, Huber W, Haering CH, Mirny L, and Spitz F. 2017. Two independent modes of chromatin organization revealed by cohesin removal. Nature. 551(7678):51–56.

104. Schule B, Oviedo A, Johnston K, Pai S, and Francke U. 2005. Inactivating mutations in ESCO2 cause SC phocomelia and Roberts syndrome: no phenotype-genotype correlation. Am J Hum Genet. 77:1117–1128.

105. Selicorni A, Mariani M, Lettieri A, and Massa V. 2021. Cornelia de Lange Syndrome: From a Disease to a Broader Spectrum. Genes (Basel). 12(7):1075.

106. Seitan VC, Banks P, Laval S, Majid NA, Dorsett D, Rana A, Smith J, Bateman A, Krpic S, Hostert A, Rollins RA, Erdjument-Bromage H, Tempst P, Benard CY, Hekimi S, Newbury SF, and Strachan T. 2006. Metazoan Scc4 homologs link sister chromatid cohesion to cell and axon migration guidance. PLoS Biol. 4(8):e242.

107. Seoane AI, and Morgan DO. 2017. Firing of Replication Origins Frees Dbf4-Cdc7 to Target Eco1 for Destruction. Curr Biol. 27(18):2849–2855.

108. Shemesh K, Sebesta M, Pacesa M, Sau S, Bronstein A, Parnas O, Liefshitz B, Venclovas C, Krejci L, and Kupiec M. 2017. A structure-function analysis of the yeast Elg1 protein reveals the importance of PCNA unloading in genome stability maintenance. Nucleic Acids Res. 45(6):3189–3203.

109. Shen D, and Skibbens RV. 2017. Temperature-dependent regulation of rDNA condensation in Saccharomyces cerevisiae. Cell Cycle. 16(11):1118–1127.

110. Shen D, and Skibbens RV. 2020. Promotion of Hyperthermic-Induced rDNA Hypercondensation in Saccharomyces cerevisiae. Genetics. 214(3):589–604.

111. Shi D, Zhao S, Zuo MQ, Zhang J, Hou W, Dong MQ, Cao Q, and Lou H. 2020. The acetyltransferase Eco1 elicits cohesin dimerization during S phase. J Biol Chem. 295(22):7554–7565.

112. Shi Z, Gao H, Bai XC, and Yu H. 2020. Cryo-EM structure of thew human cohesin-NIPBL-DNA complex. Science. 368(6498):1454–1459.

113. Sikorski RS, and Hieter P. 1989. A system of shuttle vectors and yeast host strains designed for efficient manipulation of DNA in Saccharomyces cerevisiae. Genetics. 122(1):19–27.

114. Sjögren C, and Nasmyth K. 2001. Sister chromatid cohesion is required for postreplicative double-strand break repair in Saccharomyces cerevisiae. Curr Biol. 11(12):991–995.

115. Skibbens RV, Corson LB, Koshland D, and Hieter P. 1999. Ctf7p is essential for sister chromatid cohesion and links mitotic chromosome structure to the DNA replication machinery. Genes Dev. 13(3):307–319.

116. Skibbens RV. 2019. Condensins and cohesins - one of these things is not like the other! J Cell Sci. 132(3):jcs220491.

117. Smithells RW, and Newman CG. 1992. Recognition of thalidomide defects. J. Med. Genet. 29:716–723.

118. Song J, Lafont L, Chen J, Wu FM, Shirahige K, and Rankin S. 2012. Cohesin acetylation promotes sister chromatid cohesion only in association with the replication machinery. J Biol Chem. 287(41):34325–34336.

119. Srinivasan M, Scheinost JC, Petela NJ, Gligoris TG, Wissler M, Ogushi S, Collier JE, Voulgaris M, Kurze A, Chan KL, Hu B, Costanzo V, and Nasmyth KA. 2018. The Cohesin Ring Uses Its Hinge to Organize DNA Using Non-topological as well as Topological Mechanisms. Cell. 173(6):1508–1519.

120. Stead K, Aguilar C, Hartman T, Drexel M, Meluh P, and Guacci V. 2003. Pds5p regulates the maintenance of sister chromatid cohesion and is sumoylated to promote the dissolution of cohesion. J Cell Biol. 163(4):729–741.

121. Ström L, Lindroos HB, Shirahige K, and Sjögren C. 2004. Postreplicative recruitment of cohesin to double-strand breaks is required for DNA repair. Mol Cell. 16(6):1003–1015.

122. Ström L, Karlsson C, Lindroos HB, Wedahl S, Katou Y, Shirahige K, and Sjögren C. 2007. Postreplicative formation of cohesion is required for repair and induced by a single DNA break. Science. 317(5835):242–245.

123. Sullivan M, Higuchi T, Katis VL, and Uhlmann F. 2004. Cdc14 phosphatase induces rDNA condensation and resolves cohesin-independent cohesion during budding yeast anaphase. Cell. 117(4):471–482.

124. Sun H, Zhang J, Xin S, Jiang M, Zhang J, Li Z, Cao Q, and Lou H. 2019. Cul4-Ddb1 ubiquitin ligases facilitate DNA replication-coupled sister chromatid cohesion through regulation of cohesin acetyltransferase Esco2. PLoS Genet. 15(2):e1007685.

125. Talarek N, Gueydon E, and Schwob E. 2017. Homeostatic control of START through negative feedback between Cln3-Cdk1 and Rim15/Greatwall kinase in budding yeast. Elife. 6:e26233.

126. Tanaka K, Yonekawa T, Kawasaki Y, Kai M, Furuya K, Iwasaki M, Murakami H, Yanagida M, and Okayama H. 2000. Fission yeast Eso1p is required for establishing sister chromatid cohesion during S phase. Mol Cell Biol. 20(10):3459–3469.

127. Tedeschi A, Wutz G, Huet S, Jaritz M, Wuensche A, Schirghuber E, Davidson IF, Tang W, Cisneros DA, Bhaskara V, Nishiyama T, Vaziri A, Wutz A, Ellenberg J, and Peters JM. 2013. Wapl is an essential regulator of chromatin structure and chromosome segregation. Nature. 501(7468):564–568.

128. Teufel L, Tummler K, Flöttmann M, Herrmann A, Barkai N, and Klipp E. 2019. A transcriptome-wide analysis deciphers distinct roles of G1 cyclins in temporal organization of the yeast cell cycle. Sci Rep. 9(1):3343.

129. Thorvaldsdóttir H, Robinson JT, and Mesirov JP. 2013. Integrative Genomics Viewer (IGV): high-performance genomics data visualization and exploration. Briefings in Bioinformatics. 14:178–192.

130. Tong K, and Skibbens RV. 2014. Cohesin without cohesion: a novel role for Pds5 in Saccharomyces cerevisiae. PLoS One. 9(6):e100470.

131. Tonkin ET, Wang TJ, Lisgo S, Bamshad MJ, and Strachan T. 2004. NIPBL, encoding a homolog of fungal Scc2-type sister chromatid cohesion proteins and fly Nipped-B, is mutated in Cornelia de Lange syndrome. Nat Genet. 6(6):636–641.

132. Tóth A, Ciosk R, Uhlmann F, Galova M, Schleiffer A, and Nasmyth K. 1999. Yeast cohesin complex requires a conserved protein, Eco1p(Ctf7), to establish cohesion between sister chromatids during DNA replication. Genes Dev. 3(3):320-333.

133. Ubersax JA, Woodbury EL, Quang PN, Paraz M, Blethrow JD, Shah K, Shokat KM, and Morgan DO. 2003. Targets of the cyclin-dependent kinase Cdk1. Nature. 425(6960):859–864.

134. Uhlmann F, and Nasmyth K. 1998. Cohesion between sister chromatids must be established during DNA replication. Curr Biol. 8(20):1095–1101.

135. Unal E, Arbel-Eden A, Sattler U, Shroff R, Lichten M, Haber JE, and Koshland D. 2004. DNA damage response pathway uses histone modification to assemble a double-strand break-specific cohesin domain. Mol Cell. 16(6):991–1002.

136. Unal E, Heidinger-Pauli JM, and Koshland D. 2007. DNA double-strand breaks trigger genome-wide sister-chromatid cohesion through Eco1 (Ctf7). Science. 317(5835):245–248.

137. Unal E, Heidinger-Pauli JM, Kim W, Guacci V, Onn I, Gygi SP, and Koshland DE. 2008. A molecular determinant for the establishment of sister chromatid cohesion. Science. 321(5888):566–569.

138. Vega H, Waisfisz Q, Gordillo M, Sakai N, Yanagihara I, Yamada M, van Gosliga D, Kayserili H, Xu C, Ozono K, et al. 2005. Roberts syndrome is caused by mutations in ESCO2, a human homolog of yeast ECO1 that is essential for the establishment of sister chromatid cohesion. Nat Genet. 37:468–470.

139. Vega H, Gordillo M, and Jabs EW. 2020. ESCO2 Spectrum Disorder. In: Adam MP, Ardinger HH, Pagon RA, Wallace SE, Bean LJH, Gripp KW, Mirzaa GM, Amemiya A, editors. GeneReviews® [Internet]. Seattle (WA): University of Washington, Seattle; 1993–2022.

140. Whelan G, Kreidl E, Wutz G, Egner A, Peters JM, and Eichele G. 2012. Cohesin acetyltransferase Esco2 is a cell viability factor and is required for cohesion in pericentric heterochromatin. EMBO J. 31(1):71–82.

141. Williams BC, Garrett-Engele CM, Li Z, Williams EV, Rosenman ED, and Goldberg ML. 2003. Two putative acetyltransferases, san and deco, are required for establishing sister chromatid cohesion in Drosophila. Curr Biol. 13(23):2025–2036.

142. Woodman J, Hoffman M, Dzieciatkowska M, Hansen KC, and Megee PC. 2015. Phosphorylation of the Scc2 cohesin deposition complex subunit regulates chromosome condensation through cohesin integrity. Mol Biol Cell. 26(21):3754–3767.

143. Wu PS, Enervald E, Joelsson A, Palmberg C, Rutishauser D, Hällberg BM, and Ström L. 2020. Post-translational Regulation of DNA Polymerase eta, a Connection to Damage-Induced Cohesion in Saccharomyces cerevisiae. Genetics. 216(4):1009–1022.

144. Wutz G, Várnai C, Nagasaka K, Cisneros DA, Stocsits RR, Tang W, Schoenfelder S, Jessberger G, Muhar M, Hossain MJ, Walther N, Koch B, Kueblbeck M, Ellenberg J, Zuber J, Fraser P, and Peters JM. 2017. Topologically associating domains and chromatin loops depend on cohesin and are regulated by CTCF, WAPL, and PDS5 proteins. EMBO J. 36(24):3573-3599.

145. Xiang S, and Koshland D. 2021. Cohesin architecture and clustering in vivo. Elife. 10:e62243.

146. Zhang N, Kuznetsov SG, Sharan SK, Li K, Rao PH, and Pati D. 2008. A handcuff model for the cohesin complex. J Cell Biol. 183(6):1019–1031.

147. Zhang J, Shi X, Li Y, Kim BJ, Jia J, Huang Z, Yang T, Fu X, Jung SY, Wang Y, Zhang P, Kim ST, Pan X, and Qin J. 2008. Acetylation of Smc3 by Eco1 is required for S phase sister chromatid cohesion in both human and yeast. Mol Cell. 31(1):143–151.

148. Zhang J, Shi D, Li X, Ding L, Tang J, Liu C, Shirahige K, Cao Q, and Lou H. 2017. Rtt101-Mms1-Mms22 coordinates replication-coupled sister chromatid cohesion and nucleosome assembly. EMBO Rep. 18(8):1294–1305.

149. Zhang W, Yeung CHL, Wu L, and Yuen KWY. 2017. E3 ubiquitin ligase Bre1 couples sister chromatid cohesion establishment to DNA replication in Saccharomyces cerevisiae. Elife. 6:e28231.

150. Zuilkoski CM, and Skibbens RV. 2020a. PCNA promotes context-specific sister chromatid cohesion establishment separate from that of chromatin condensation. Cell Cycle. 19(19):2436–2450.

151. Zuilkoski CM, and Skibbens RV. 2020b. PCNA antagonizes cohesin-dependent roles in genomic stability. PLoS One. 15(10):e0235103.

